# Caspase prime-side active-site characterization with non-hydrolyzable peptides assists in the design of a caspase-7-selective irreversible probe

**DOI:** 10.1101/2024.01.30.578085

**Authors:** Angelo Solania, Janice H. Xu, Dennis W. Wolan

## Abstract

Limited structural information and biochemical studies are available that demonstrate how the prime side of cysteine protease active sites, such as for human caspases, are used for substrate recognition and how these binding regions can be exploited in the design of inhibitors and probes. Reported small molecules that interact with, and are optimized for, the prime side of caspase active sites are limited to methylketone leaving groups and other nonpeptidic inhibitor moieties, such as aza-Michael acceptors. We present the design, synthesis, and co-complex x-ray structures of the first prime-side elongated non-hydrolyzable peptidomimetic ketomethylene inhibitors designed to interrogate the S4-S4’ active-site binding pockets of the executioner caspases-3 and -7. In addition to our structures depicting the first examples of an active-site cysteine in complex with a P1 residue trapped in a non-covalent tetrahedral intermediate, we elucidated prime-side binding interactions for pockets S1’ through S4’ with our biologically relevant peptide Ac-DEVD-Propionate-AAA. Despite the substantial homology among the caspase active sites, we identified a key difference in the prime-side architecture within binding distance to the P2’ inhibitor alanine whereby caspase-3 F128 is substituted for caspase-7 Y151. We exploited this prime-side difference in side chains and their reactivities in the design of non-hydrolyzable ketomethylene-based probes bearing a C-terminal tyrosine-reactive 4-phenyl-1,2,4-triazole-3,5-dione moiety. Our probe selectively labels caspase-7 over caspase-3 and we posit that further characterization of protease active-site prime sides with similar non-hydrolyzable molecules will yield additional tool-like compounds that will assist in establishing the non-redundant roles of caspase family members and other cysteine proteases.

---

The ability to regulate human caspases remains a formidable goal since this family of cysteine proteases was first established as a nexus in apoptotic programmed cell death almost 30 years ago^1,2^. More recently, studies have indicated that the biological roles of caspases are far more complex, as their proteolytic activities are now implicated in many non-lethal cellular processes^3-10^. Outstanding questions with respect to the individual substrates and cellular events involving caspases as well as the potential of these proteases as therapeutic targets remain to be addressed. Genetic approaches, including CRISPR-Cas9 and cellular and animal knockout studies have provided tremendous insights into the individual contributions of selected caspase family members^11-18^. However, these methodologies have limitations, including the inability to discriminate enzymatic activity from protein:protein interactions, as well as the inherent lack of temporal resolution to differentiate first-order mechanisms and secondary effects of target protein function(s) in living systems. Alternative approaches to genetic regulation include the application of mechanism-based caspase peptidyl small-molecule inhibitors with C-terminal activated methyl ketones for covalent attachment. While the chemical-based regulation of caspase activity provides a complementary set of techniques capable of dose-dependent experiments with resolvable kinetic and spatial resolution, molecules require extreme specificity to confidently attribute cellular responses to individual proteases. Such selectivity is difficult to obtain due to the high structural homology and overlapping substrate specificities across the caspases.

Irreversible inhibitors containing C-terminal warheads such as halomethylketones (*i*.*e*., CMK, chloromethylketone; FMK, fluoromethylketone), vinylsulfones, epoxides, diisopropylfluorophosphate (DFP) and acyloxymethylketones (AOMK) have been indispensable to establishing the specificity and catalytic mechanisms of proteases^19-21^. These molecules pair an electrophilic ketone at the P1 position with a leaving group (*e*.*g*., CMK) on the adjacent prime-side carbon and this warhead typically replaces the entire prime-side recognition motif of the natural peptide substrate (*e*.*g*., Gly-Ala-Ala-Ala from Ac-DEVD-GAAA-OH) (Figure 1A). The leaving group is displaced upon nucleophilic attack by the active-site cysteine (or histidine of serine proteases) to form an irreversible, stable bond. Such covalent substrate mimetics have provided critical insights into the structural characterization of protease active sites, as best exemplified by the first co-complex structure of papain bound with a CMK-containing peptidomimetic over 40 years ago^22^. However, due to the replacement of prime-side residues, the expansive collection of small molecules and peptidomimetics that selectively target individual proteases, including the caspases, have almost exclusively focused on the optimization of interactions within the non-prime side of active sites^23-28^.

**Figure 1.**
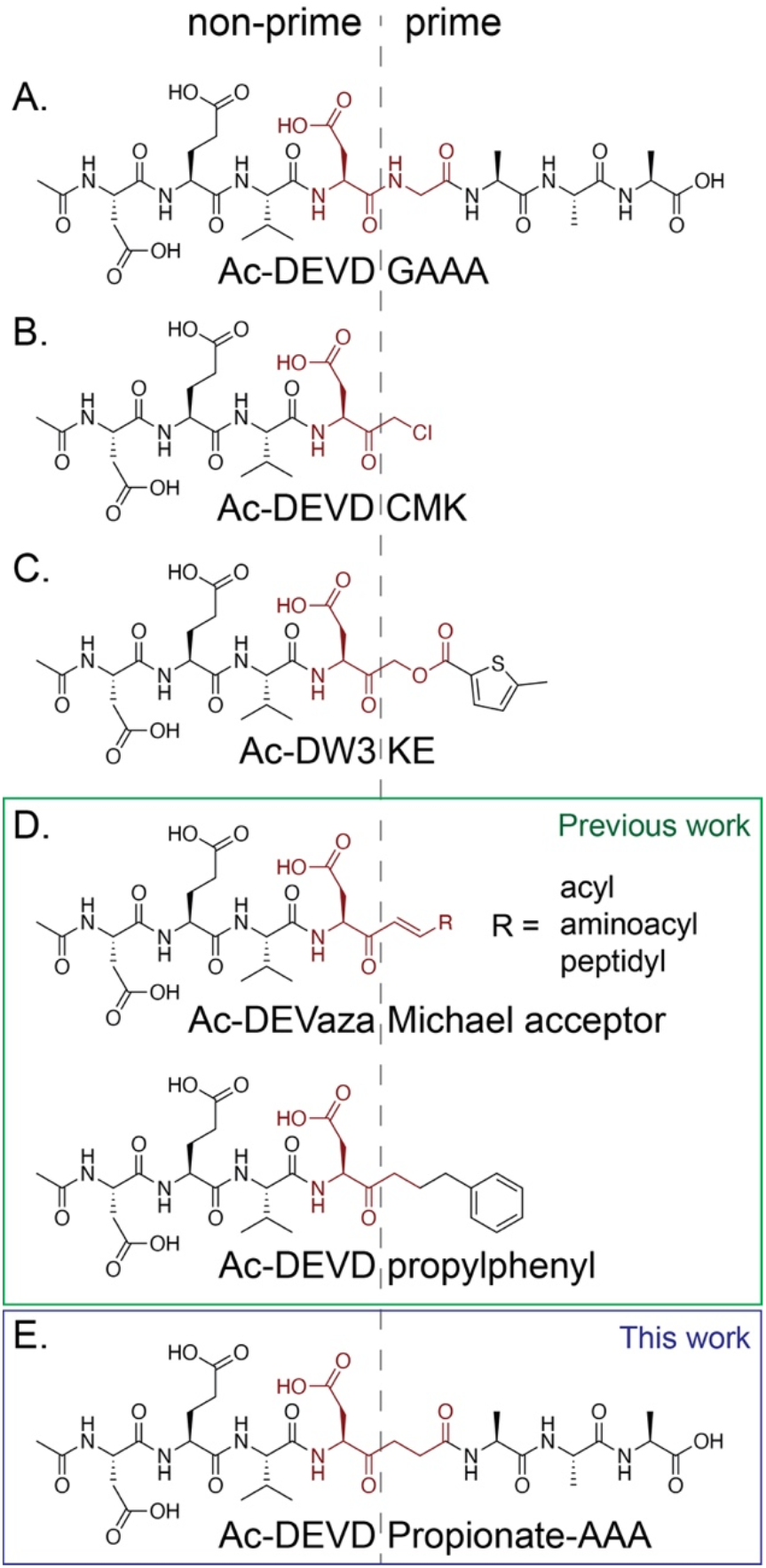
Peptide-based caspase inhibitors and a new non-hydrolyzable prime-side 8-mer peptide. (A) Schematic of an 8-mer peptide substrate spanning the scissile bond with four residues on the prime side (separated by dotted line). (B,C) Available inhibitors include: pan-caspase-activated chloromethylketone Ac-DEVD-CMK and a casp-3-selective acyloxymethylketone Ac-DW3-KE. (D) Structures of compounds with active site prime-side interactions previously used to structurally characterize the prime subsites of caspases^27,29^. (E) Our non-hydrolyzable 8-mer peptidomimetic ketomethylene inhibitor Ac-DEVD-Propionate-AAA designed to establish the biologically relevant interactions with the prime side of the caspase active site.

The limited number of chemical tools that incorporate prime-side recognition motifs has resulted in a remarkable disparity between the understanding garnered from non-prime and prime-side structural characterizations with substrate mimetics. This lack of prime-side information is attributable to the inherent characteristics of currently available molecules, such as the activated methylketone inhibitors (*i*.*e*., CMK, AOMK) (Figures 1B and 1C). Despite these limitations, structural characterization of the prime-side binding region of caspases has been reported using aza-Michael acceptors and ketomethylene inhibitors^27,29^. These molecules provide some structural insights on the immediate prime-side binding pocket relative to the scissile bond. Caution is warranted in the interpretation of these structural data, as all the reported molecules differ substantially from the biologically relevant peptide scaffold (Figure 1D). Collectively, studies based on these inhibitors suggest that the prime side of the caspase active site has limited significance to the protease’s specificity. However, we show with the application and optimization of non-hydrolyzable peptides that prime-side residues can be exploited to generate caspase-selective probes.

We determined a high-resolution x-ray crystal structure of human caspase-8 (casp-8) in complex with an AOMK peptidyl inhibitor Ac-DW3-5-methyl-2-thiophene carboxylic acid (KE). Surprisingly, the structure featured a tetrahedral transition-state (TS) intermediate between the active-site cysteine side chain and P1 carbonyl of the bound inhibitor. While structural studies of serine proteases and their respective probes have established and verified tetrahedral transition-state intermediates^30-35^, this is the first high-resolution structural evidence of a tetrahedral intermediate in the active site of a cysteine protease with an AOMK. We replicated this TS intermediate in the structures of casp-3 and casp-7 using peptidyl ketomethylene inhibitors. These non-hydrolyzable peptides, including a proof-of-concept ketomethylene inhibitor, Ac-DEVD-Propionate, and a prime-side elongated substrate mimetic, Ac-DEVD-Propionate-AAA, were prepared via solid phase using an Fmoc-protected dipeptidomimetic building block, Fmoc-Asp(OBzl)-Propionate (Figure 1E). Our collection of human caspase crystal structures bound with a new class of biologically relevant, uncleavable caspase-directed peptidomimetics represents the first three-dimensional characterization of the prime sides of casp-3 and casp-7. Ultimately, our non-hydrolyzable peptides and crystal structures guided the design of an irreversible casp-7-selective probe that represents the first small molecule capable of differentially detecting and labeling active casp-7 from the highly homologous casp-3.

## Results and Discussion

### Ac-DW3-KE forms a tetrahedral intermediate in complex with caspase-8

We determined a 1.48 Å co-crystal structure of casp-8 in complex with Ac-DW3-KE to establish how the inhibitor is selective for casp-3 from a structural perspective^36^. The active-site electron density revealed the formation of an unexpected adduct between Ac-DW3-KE and the catalytic cysteine C360. Unlike other caspase co-crystal structures featuring activated methylketones, strong naïve density was observed in the S1’ and S2’ subsite of casp-8 (Figure S1). We hypothesized that this density corresponded to the 5-methyl-2-thiophene carboxylate of Ac-DW3-KE that is expected to be displaced after alkylation of the active site cysteine. The KE leaving group fit the density and the thiophene sulfur positioned within strongly diffracting electron density indicative of a sulfur, suggesting the inhibitor is bound in the unreacted state (Figure S1). Moreover, the density shows that the side chain of C360 binds to the P1 carbonyl of Ac-DW3-KE (Figure 2A), as opposed to the adjacent α’ methylene observed with the previously reported co-crystal structures of Ac-DW3-KE in complex with casp-3 (PDB ID 6CKZ) and casp-7 (PDB ID 6CL1)^36^. Closer inspection of the angles and distances surrounding the P1 carbonyl of Ac-DW3-KE further supports that the adduct formed within the caspase-8 active site adopts a tetrahedral geometry (Figures 2B and 2C). We posited that replacement of the leaving group to generate a ketomethylene inhibitor would permit the design of non-hydrolyzable substrate mimetics that would mimic the tetrahedral conformation of the P1 carbonyl as well as afford structural characterization of the prime side of caspase active sites.

**Figure 2.**
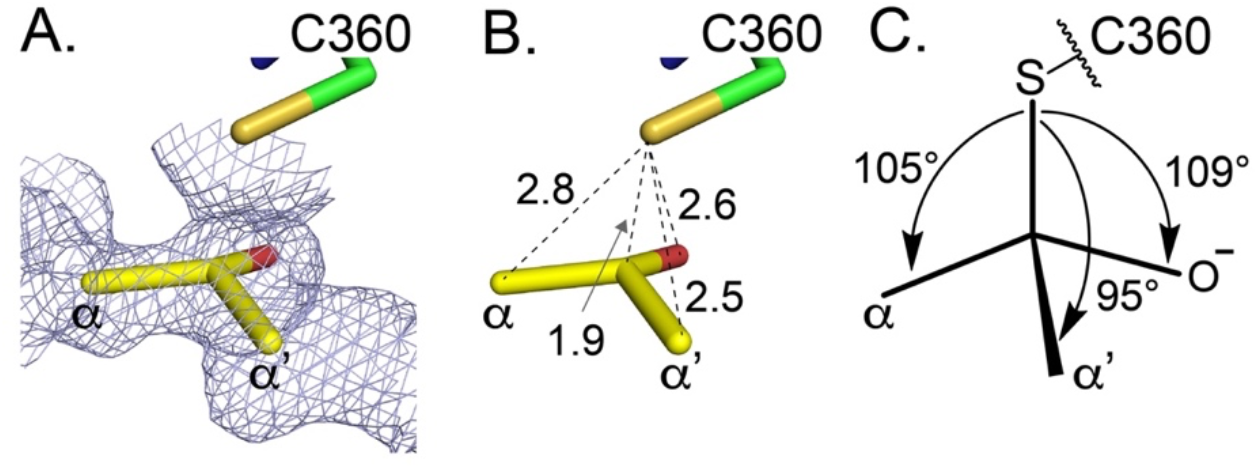
Ac-DW3-KE adopts a tetrahedral intermediate in the caspase-8 co-complex crystal structure. (A) Zoomed view of the naïve *f*_*o*_-*f*_*c*_ density of the Ac-DW3-KE carboxylate adduct (yellow carbon, red oxygen) with casp-8 C360 (green carbon, mustard sulfur) contoured at 1σ. Residues P4-P2 and the ketomethylene portion of Ac-DW3-KE were removed for clarity. (B) Bond distances between C360 sulfur and the four nearest atoms of Ac-DW3-KE suggest casp-8 binds to P1 carbonyl carbon of ketomethylene inhibitor. (C) Angles formed between C360 sulfur and Ac-DW3-KE carbon 1 substituents adopt the nearly ideal 109.5° for tetrahedral geometry.

### Synthesis of ketomethylene inhibitor Ac-DEVD-Propionate

Design of the non-hydrolyzable peptide began with the synthesis of a dipeptidomimetic building block, Fmoc-Asp(OBzl)-Propionate that would be amenable to standard solid-phase peptide synthesis (SPPS) and incorporation into any canonical caspase-specific peptide sequence. While the synthesis of several different ketomethylene building blocks suitable for peptide synthesis have been reported^37-40^, the syntheses typically require many steps and are narrow in substrate scope. We devised a combination of these synthetic routes to access the requisite Fmoc-protected dipeptide mimetic building block, Fmoc-Asp(OBzl)-Propionate (Scheme 1). Briefly, the hydroxyl group of N-Boc-*L*-homoserine was protected and the carboxylate converted into a Weinrab amide before treatment with butenyl magnesium bromide. The hydroxyl was deprotected, oxidized, and the OBzl group was protected prior to oxidation of the olefin. Subsequent Boc deprotection and Fmoc-protection of the α-amine yielded the requisite dipeptide mimetic building block Fmoc-Asp(OtBu)-Propionate (please see Supporting Information for details). As previously reported, this building block was synthesized partially racemic at the P1 aspartate.

The partially racemic Fmoc-Asp(OBzl)-Propionate was used to synthesize the protected peptide Ac-D(OtBu)E(OtBu)VD(OBzl)-Propionate via SPPS (Scheme 2). The peptide was synthesized using established SPPS and hydrolyzed from the solid support to yield a mixture of Ac-DEV(*D*/*L*)D(OBzl)-Propionate isomers. The benzyl ester was left intact, as we found the presence of the protecting group substantially improved the separation resolution by achiral HPLC. The individual pure isomers were then treated with 10% Pd/C under hydrogen, filtered, and purified by reverse-phase HPLC to generate the fully deprotected peptides.

### Ac-DEVD-Propionate is a potent reversible inhibitor of caspases-3 and -7

Both diastereomers of the two fully deprotected peptides were verified by high-resolution LC-MS and subjected to inhibition studies against casp-3 to help identify the correct *L* stereoisomer, as the active site prefers *L* over *D* P1 aspartate side chains. Casp-3 preincubated with increasing concentrations of the either peptide was subjected to a fluorogenic activity assay using Ac-DEVD-AMAC substrate cleavage. As predicted, one of the diastereomers had substantially higher inhibition potential over the other, indicated by an IC_50_ of 4.75 ± 0.60 nM compared to IC_50_ of 232 ± 26 nM (Figure 3A). With the correct canonical (*L*) P1 aspartate-containing peptide in hand, we next measured the inhibition potential of Ac-DEVD-Propionate against casp-7 and casp-8.

**Figure 3.**
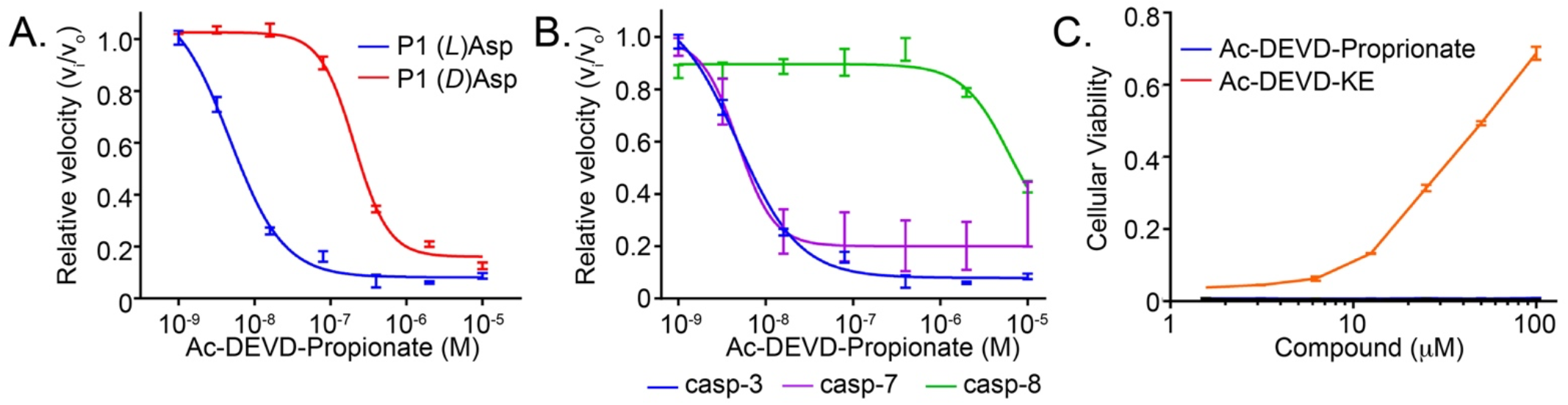
*In vitro* and cell-based validation of ketomethylene inhibitor Ac-DEVD-Propionate. (A) Activity dose response curves of 1 nM casp-3 pre-treated with increasing concentrations of Ac-DEV(*L*)D-Propionate or Ac-DEV(*D*)Asp-Propionate shows a 10-fold reduction in IC_50_ for the (*D*)Asp enantiomer (red) relative to the (*L*)Asp enantiomer (blue) in the P1 position. (B) Activity dose response curves of Ac-DEVD-Propionate against panel of apoptotic caspases shows the peptide robustly inhibits casp-3 (blue) and casp-7 (purple), but weakly inhibits casp-8 (orange). (C) Despite the strong *in vitro* inhibitory potential of Ac-DEVD-Propionate (blue), the inhibitor did not afford protection of Jurkat cells against extrinsically induced apoptosis with 10 ng/mL MegaFasL in comparison to Ac-DEVD-KE (blue). Cellular viability was measured using Cell-TiterGlo®.

Similar dose-dependent activity assays against casp-7 and casp-8 preincubated with Ac-DEVD-Propionate showed that the peptide only effectively inhibited casp-7. To our surprise, casp-8 activity was only reduced at high concentrations of Ac-DEVD-Propionate despite the ability of casp-8 to efficiently recognize and bind DEVD-based substrates and inhibitors (Figure 3B)^23^. To determine if this lack of inhibition was due to the non-prime interactions with DEVD, the propionate, or a combination of both regions of the peptide, we synthesized a ketomethylene inhibitor bearing the preferred casp-8 recognition sequence, Ac-IETD-Propionate. Our results show that the IETD-based inhibitor was no more effective than Ac-DEVD-Propionate toward casp-8 and suggests that the non-prime canonical recognition sequence is not a contributing factor to the decreased potency. We hypothesize that the poor inhibition is likely attributable to casp-8 forming a less stable adduct with the ketomethylene electrophile relative to casp-3 and casp-7. Of note, Ac-IETD-Propionate did not inhibit casp-3 and casp-7 at the concentrations tested (Figure S2).

We next assessed the ability of Ac-DEVD-Propionate to protect Jurkat T lymphocytes from extrinsically induced apoptosis. Jurkat cells were preincubated with increasing concentrations of Ac-DEVD-Propionate, Ac-DEVD-KE (positive control), or vehicle (negative control, DMSO) for 30 min prior to introduction of MegaFasL. After 4 h, apoptosis was measured by Cell-TiterGlo®. Despite the ability of Ac-DEVD-Propionate to effectively inhibit casp-3 and casp-7 activity under *in vitro* conditions, the compound was incapable of protecting against apoptosis in comparison to Ac-DEVD-KE (Figure 3C). We posit that the reversible binding of Ac-DEVD-Propionate compared to the irreversible inhibition of caspase activity by Ac-DEVD-KE is responsible for the inability to protect Jurkat cells from MegaFasL-induced apoptosis.

### Ac-DEVD-Propionate binds to caspases-3 and -7 with a tetrahedral P1 carbonyl

We determined the co-complex crystal structures of casp-3 and casp-7 bound with Ac-DEVD-Propionate to 1.50 and 2.60 Å resolution, respectively, to eluicidate the prime-side active-site interactions with the propionate moeity. Similar to the casp-8:Ac-DW3-KE structure, naïve electron density was clearly present in the S1’ subsite of both casp-3 and casp-7 structures and the propionate group was readily modeled into the density with B factors similar to the surrounding active site residues (Figure S3). Furthermore, the electron density depicted the casp-3 active-site cysteine side chain (C163) to reside in very close proximity to the P1 carbonyl carbon of Ac-DEVD-Propionate, as well as formed a tetrahedral intermediate with angles between 103-113° (Figure S3). Due to the lower resolution of the casp-7:Ac-DEVD-Propionate co-crystal structure, we cannot adequately surmise the same conclusions with as much confidence; however, the Ac-DEVD-Propionate models into the casp-7 active site similarly to casp-3 (Figure S3). Despite repeated attempts at crystallization of casp-8 in complex with Ac-DEVD- and Ac-IETD-Propionate, crystals did not diffract to reasonable resolution and we therefore cannot address why addition of the propionate group is not effective at casp-8 inhibition.

### Structures of caspases with Ac-DEVD-Propionate-AAA reveal prime-side interactions

We next extended the prime side of the inhibitor with three alanine residues by SPPS to generate Ac-DEVD-Propionate-AAA (Figure 1E). Caspases-3 and -7 readily crystallized in the presence of Ac-DEVD-Propionate-AAA and the structures were determined to 2.17 and 2.45 Å resolution, respectively. Similar to the Ac-DEVD-Propionate orientation within the caspase active site, a conserved tetrahedral geometry was observed in the density for the P1 carbonyl of Ac-DEVD-Propionate-AAA with the active site C163 of casp-3 and casp-7 positioned near the P1 carbonyl carbon. Most significant was the clear and continuous electron density for the P1’ through P4’ alanines in the casp-3 complex and through P3’ for the casp-7 structure (Figure S4). Our structures support previous studies demonstrating that only small aliphatic residues can be accomodated into the P1’ pocket as both α hydrogens of the propionate moiety are directed toward the protein (Figures 4A and 4B). Overlay of Ac-DEVD-Propionate and Ac-DEVD-Propionate-AAA bound to casp-3 show the inhibitors superimpose identically between the P4 to P1’ residues and suggests that binding to the prime-side of the protease does not influence interactions or promote conformational changes on the non-prime side (Figure 4C).

**Figure 4.**
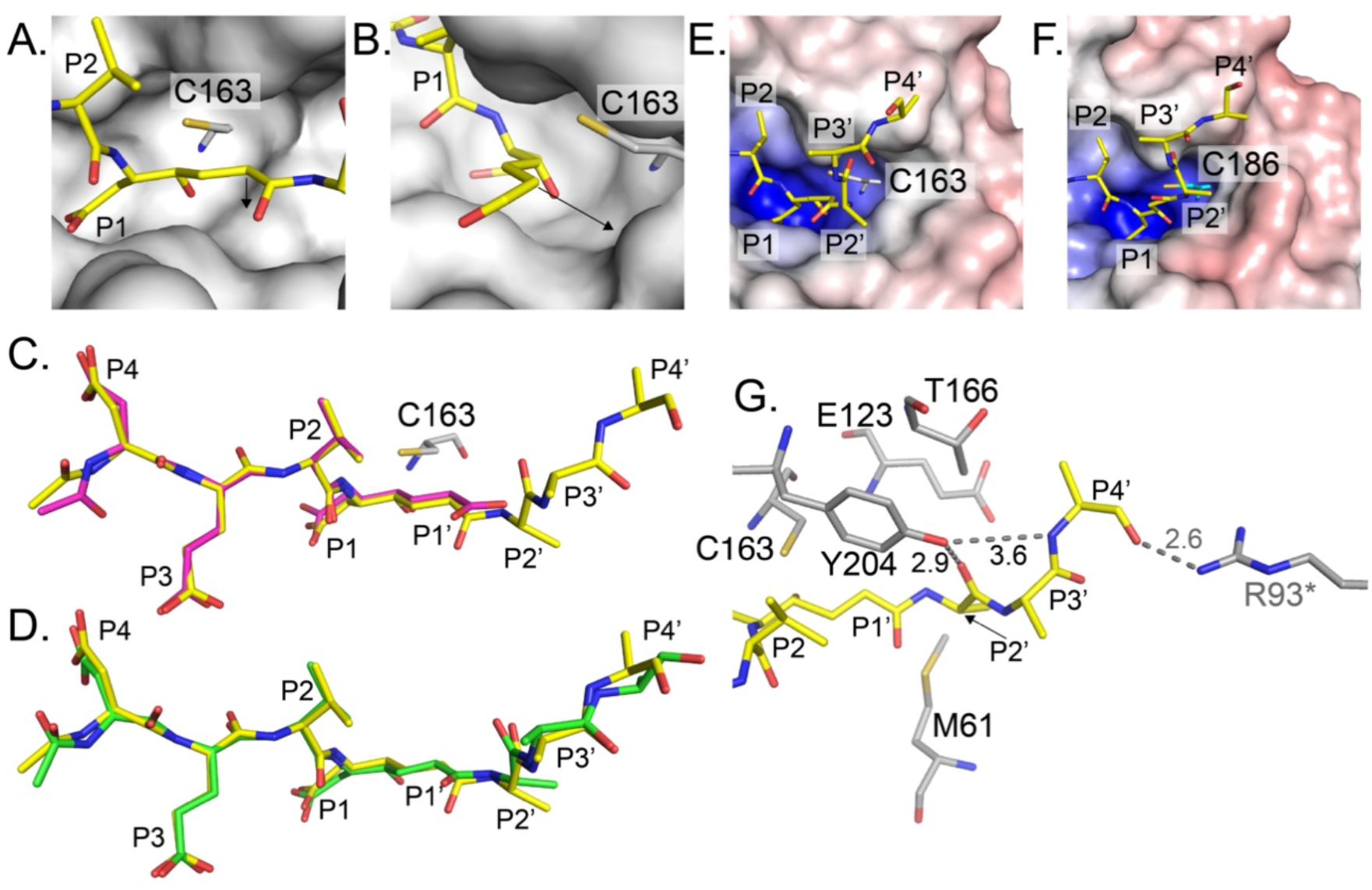
Expanded prime-side caspase inhibitor show key interactions with casp-3 and casp-7 active site pockets. (A) Casp-3:Ac-DEVD-Propionate-AAA (gray and yellow carbon, respectively) co-crystal structure highlighting the orientation of the P1’ hydrogens and insufficient space to accommodate large side chains (blue nitrogen, red oxygen, mustard sulfur). The black arrow highlights orientation of relevant stereotopic hydrogen. (B) 90° rotation of (A) with Ac-DEVD-Propionate-AAA residues P2’-P4’ removed for clarity. (C) Overlay of Ac-DEVD-Propionate (magenta) and Ac-DEVD-Propionate-AAA (yellow) bound to casp-3 (gray). (D) Overlay of Ac-DEVD-Propionate-AAA to casp-3 (yellow) and casp-7 (salmon) showing deviations of Ac-DEVD-Propionate outside of P4 to P2’. (E, F) Electrostatic potential surface of casp-3 and casp-7 prime side with Ac-DEVD-Prop-AAA bound highlighting hydrophobic environments past the P2’. Arrows shown denote the relative directions of the P2’-P4’ side chains. Blue, positive potential (≥10 mV); white, neutral potential (0 mV); red, negative potential (≤−10 mV). (G) Schematic of Ac-DEVD-Propionate-AAA in complex with casp-3 highlighting prime-side interactions and residues within 4 Å of the ligand that could provide potential interactions with expanded prime-side side chains (colored as in A). * and gray distance label denotes a non-biologically relevant crystal contact to the Ac-DEVD-Propionate-AAA P4’ main-chain carbonyl.

Caspases-3 and -7 are highly homologous in sequence and structure and the active site, spanning the protease’s surface from S4 to S4’, is over 75% identical. With respect to the non-prime side, 13 of the 17 residues within 4 Å of the bound Ac-DEVD-portion of the inhibitors (*i*.*e*., Propionate and Propionate-AAA) are identical (Figure S4). Residues that differ between the two proteases include (*e*.*g*., casp-3/casp-7) Q250/F276, N208/S234, and S209/P235 that form distal interactions at, and beyond, the P4 position. The last residue (S63/T86) appears to form van der Waals interactions with the Ac-DEVD, as it is 4 Å away from the carboxylate of the P3 glutamate. The primary difference on the non-prime side is the orientation of the P5 residue that is likely attributable to the ability of S209 in casp-3 to form a potential hydrogen bond, while the proline (P235) in casp-7 is incapable of providing a hydrogen bond.

The conformations of the Ac-DEVD-Propionate-AAA prime-side residues are superimposable among the casp-3 and -7 active sites with some minor differences in oriention of the P3’ and P4’ residues (Figure 4D). Dissimilar to the non-prime-side of the active site with pockets for peptidyl substrate side chains (P4-P1), the prime side of both casp-3 and casp-7 lack clearly defined surfaces to accommodate and bind side chains and consists of limited electrostatic surface potential (Figures 4E and 4F). One of the most sigificant contributors to interactions on the prime side includes the casp-3 Y204 side chain (Y230 in casp-7) which can provide potential hydrogen bonds to both the main-chain carbonyl oxygen of P2’ and the main-chain amide nitrogen of P3’ (Figure 4G). This hydrogen-bond network orients and stabilizes the P2’ and P3’ side chains such that the P2’ side chain is directed toward the protein surface and the P3’ side chain appears to be oriented away from the active site and is solvent exposed (Figures 4E and 4F). Of note, the measured distance between the Ac-DEVD-Propionate-AAA P3’ main-chain amide nitrogen and Y230 side chain in the casp-7 is 4.6 Å and suggests a limited contribution to the substrate orientation in comparison to casp-3. M61 and E123 (M84 and E146 in casp-7) are poised within the casp-3 prime side to provide potential interations with larger P2’ substrate side chains and T166 (T189 in casp-7) can provide potential hydrogen bonds with an extended P4’ substrate side chain (Figures 4E, 4F, and S4). Importantly, accomodating larger P2’ and P4’ side chains on peptide substrates may influence the binding and orientation of the substrate prime-side main chain and is particularly relevant for P3’ and P4’ as limited main-chain interactions occur with the casp-3 and casp-7 active sites (Figure 4G). The quality of electron density diminishes and the B factors increase as the peptide extends into the P4’ region for both the casp-3 and casp-7 co-complex structures and suggests that the inhibitor becomes increasingly flexible. However, the P4’ main-chain carbonyl in the casp-3 structure has definable density relative to the casp-7 structure and is likely attributable to both structure resolution (2.17 Å for casp-3 and 2.45 Å for casp-7, see Supporting Information for structure statistics), as well as a potential hydrogen bond provided by R93 from a crystal contact absent in casp-7 crystal packing (Figures 4G and S4). While this non-biologically relevant hydrogen bond may stabilize the P4’ main-chain carbonyl, this interaction likely does not influence the orientation and conformation of the Ac-DEVD-Propionate-AAA prime-side residues.

Our casp-3 and -7 structures bound with Ac-DEVD-Propionate and Ac-DEVD-Propionate-AAA support previous reports primarily from proteomics analyses that caspases prefer substrates with small side chains in the P1’ position^26,28,41-44^. Furthermore, our structural analyses support observations that incorporation of hydrophilic moieties in the prime side can maintain and, in some cases, improve the potency of active-site directed molecules for caspases.

### A caspase-7-selective irreversible probe employing prime-side structural information

Beyond the immediate cluster of caspase residues that directly interact with the prime side of Ac-DEVD-Propionate-AAA, there are several residues that we posited could be exploited in the design of a new caspase-selective inhibitor. The P2’ β-carbon of Ac-DEVD-Propionate-AAA is approximately 4.5 Å from the hydroxyl of Y151 in casp-7 and 6.1 Å from the homologous F128 in casp-3 (Figures 5A and 5B). Interestingly, the casp-7 Y151 is conserved across all human apoptotic caspases, including caspases-2, -6, -8, -9, and -10 and is a cysteine for inflammatory caspases-1, -4, and -5 (Figure 5C). Only casp-3 consists of a phenylalanine at this position. While the canonical non-prime-side tetrapeptide sequence DEVD is selective towards casp-3 and casp-7, and casp-8 based on substrate kinetics and non-prime-side-based inhibitors^25,36,45^, we hypothesized selectivity for casp-7 over casp-3 could be accomplished by incorporating a tyrosine-selective functionality at the P2’ position of the extended Ac-DEVD-Propionate-based inhibitor. Mutational modeling substituting the Ac-DEVD-Propionate-AAA P2’ Ala with a glutamate showed several common glutamate rotamers position the side-chain carboxylic acid near the Y151 side-chain hydroxyl suggesting that a reactive moiety could be added to the inhibitor that would covalently modify Y151 of casp-7 and not F128 of casp-3 (Figure 5B, inset).

**Figure 5.**
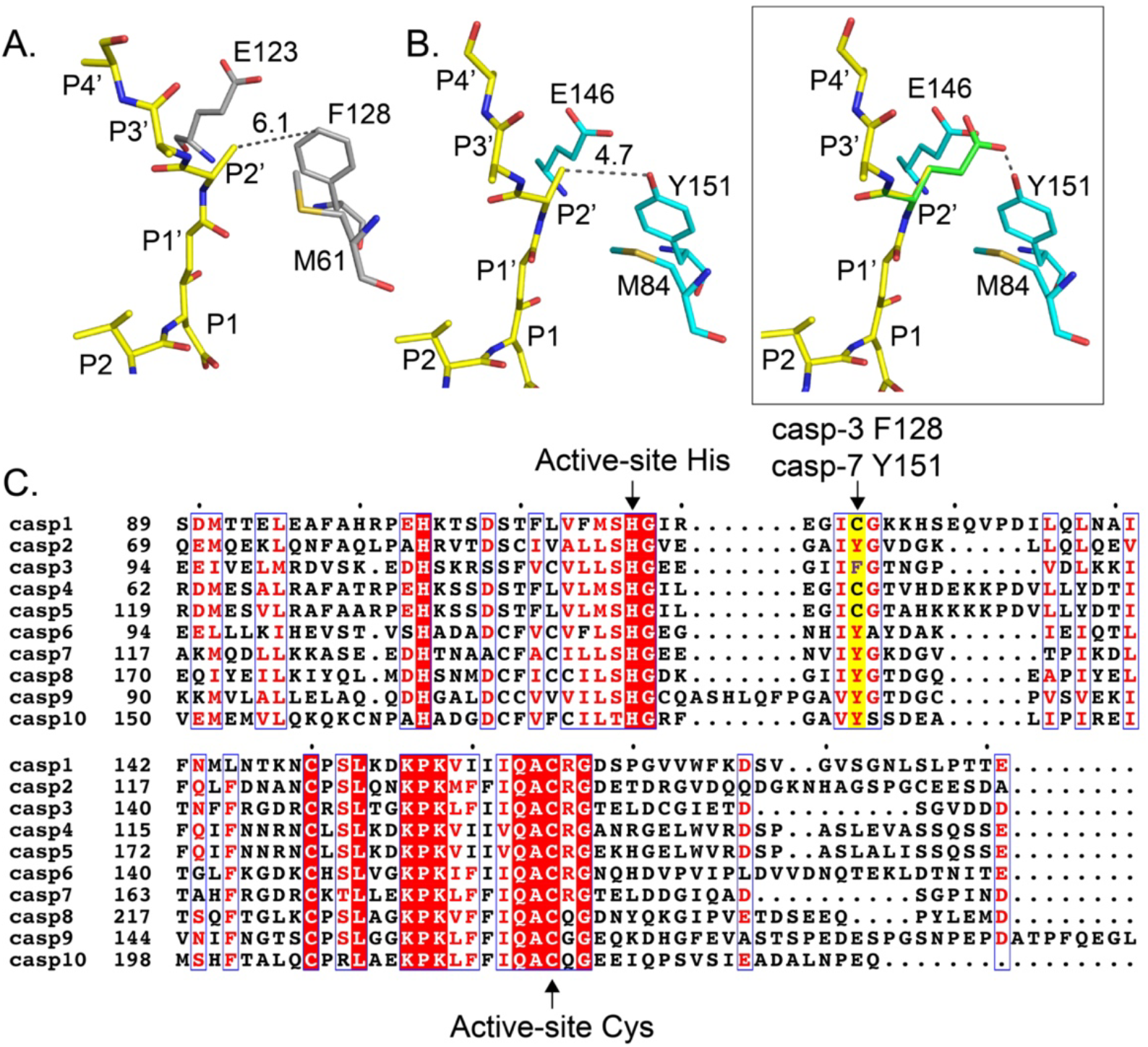
Differences in casp-3 and casp-7 prime-side regions of the active site for the design of a casp-7-selective probe. (A, B) Casp-3 (gray carbon) and casp-7 (teal carbon) prime-side active-site residues situated within 4 Å of Ac-DEVD-Propionate-AAA (yellow carbon). The P2’ Ala β-carbon is approximately 6.1 Å from casp-3 F128 and 4.7 Å from casp-7 Y151. (B, inset) PyMol-simulated mutation of Ac-DEVD-Propionate-AAA P2’ alanine to glutamate models how the side-chain carboxylic acid can adopt conformations that places the residue within range of Y151 for covalent modification (one rotamer shown). (C) Clustal Omega-based alignment^58^ and Espript-based^59^ schematic representation of human casp-1, -2, -3, -4, -5, -6, -7, -8, -9, and -10 (casp-12 and casp-14 excluded) focused on the casp-7 Y151 residue (yellow box). The F128 residue of casp-3 is colored purple and is unique across all human caspases. The catalytic histidine and cysteine residues are labeled accordingly.

We initially posited it may be possible to target Y151 by converting the Ac-DEVD-Propionate-AAA P2’ Ala side chain into Ac-DEVD-Propionate-E(chloromethylketone)-OH. Unfortunately, the requisite precursor was not found to be amenable to chemical synthesis due to spontaneous intramolecular cyclization (data not shown). Alternatively, we replaced the P2’ residue with AEBSF (aminoethylbenzylsulfonic acid) a general protease inhibitor known to inhibit serine proteases. Despite the ability to synthesize Ac-DEVD-Propionate-AEBSF, this sulfonyl fluoride was unable to covalently modify casp-7 (data not shown).

The use of 4-phenyl-1,2,4-triazole-3,5-dione (PTAD) has been recently reported to covalently and selectively modify tyrosine side chains under biocompatible conditions at room temperature^46^. In aqueous buffers, PTAD adds to the *ortho*-position relative to the side-chain hydroxyl (Figure 6A). We synthesized an aminophenyl-PTAD and conjugated the molecule to biotin-DEVD-Propionate to form biotin-DEVD-Propionate-3/4-PTAD (Figure 6B, Scheme 3). We assessed the ability of biotin-DEVD-Propionate-4-PTAD and biotin-DEVD-Propionate-3-PTAD to covalently modify casp-7 under *in vitro* conditions in comparison to casp-3. Recombinant casp-3 and casp-7 were incubated at 250 nM overnight at room temperature in caspase activity buffer with 1, 5, and 50 µM biotin-DEVD-Propionate-3/4-PTAD. The ability of PTAD-containing inhibitors to label casp-3 and -7 was measured by avidin blot. Prior to use, prechilled biotin-DEVD-Propionate-3/4-PTAD was treated with prechilled dibromantin to generate corresponding triazole active species and immediately used for labeling. Both biotin-DEVD-Propionate-4-PTAD and biotin-DEVD-Propionate-3-PTAD selectively labeled casp-7 with no observable labeling of casp-3 (Figure 6C). Moreover, biotin-DEVD-Propionate-4-PTAD labeled casp-7 more effectively than the 3-PTAD version at all concentrations tested. To verify PTAD specifically labelled casp-7 Y151, single-point mutagenesis of the residue to either phenylalanine or alanine were attempted. However, these casp-7 Y151 mutants failed to generate active, properly folded casp-7 suggesting that Y151 is essential to the structural stability of the protease and/or ability to self-activate during exogenous *E. coli* overexpression and/or throughout the purification process. Additional characterization of biotin-DEVD-Propionate-3/4-PTAD casp-7 inhibition kinetics and applicability in cellular assays are necessary to further establish the utility of this selective mode of casp-7 inhibition, as well as provide insight into the ability to develop non-hydrolyzable peptidyl PTAD-based inhibitors to other caspase family members with a conserved Tyr in the prime-side of the active site.

**Figure 6.**
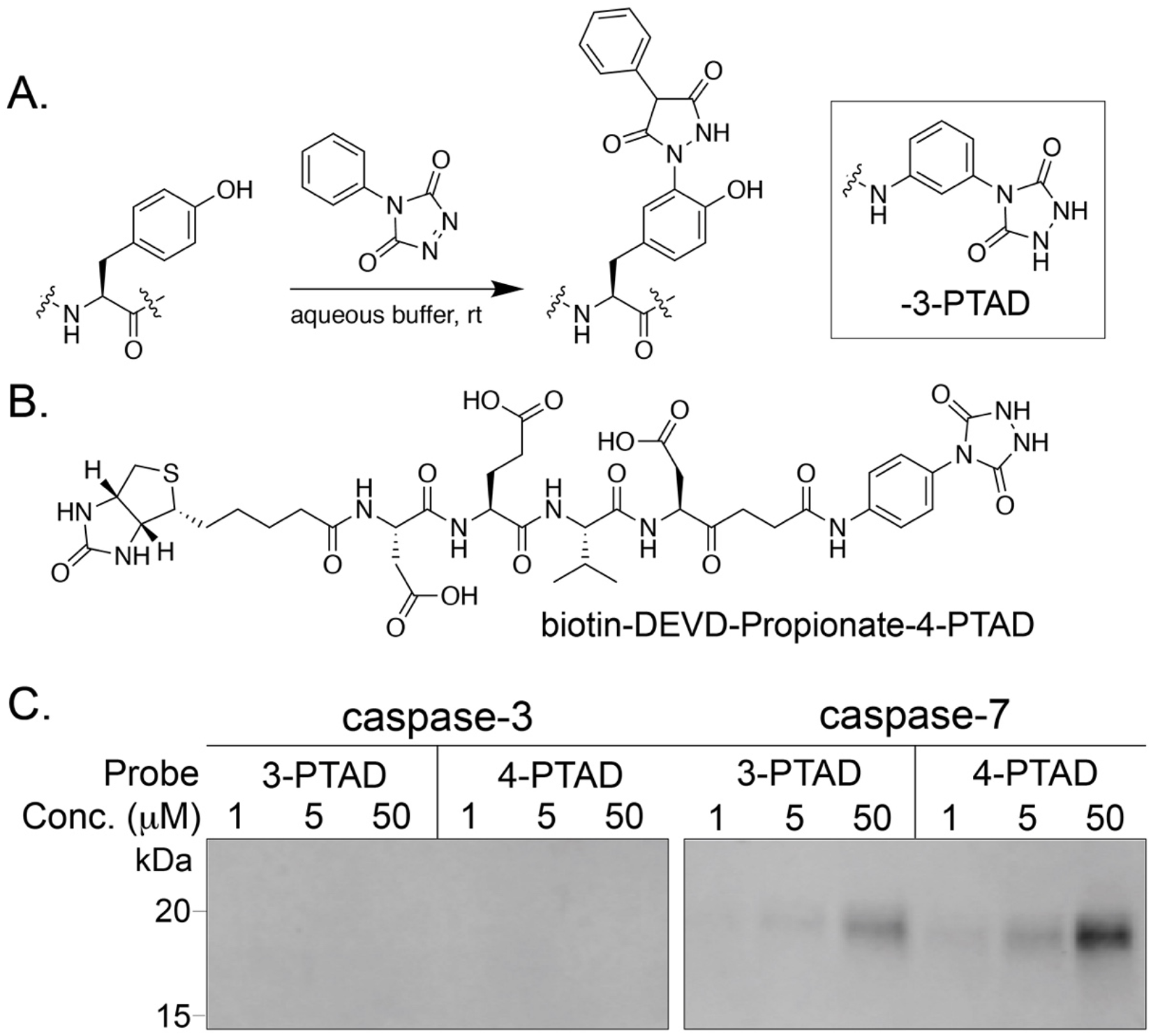
Synthesis and selective labeling of casp-7 by biotin-DEVD-Propionate-3/4-PTAD. (A) General labeling of tyrosine side chains by PTAD. (B) Chemical structure of biotin-DEVD-Propionate-4-PTAD and biotin-DEVD-Propionate-3-PTAD (inset). (C) Recombinant casp-3 and casp-7 were subjected to incubation with either biotin-DEVD-Propionate-3-PTAD or biotin-DEVD-Propionate-4-PTAD overnight and assessed for covalent labeling by avidin blot. The large domain of casp-7 (residues 24-198, 19.7 kDa) was selectively labeled by biotin-DEVD-Propionate-3/4-PTAD and no labeling of casp-3 was observed.

## Conclusions

We present the first elongated peptidyl ketomethylene inhibitors, as well as the structure of the molecules in complex with casp-3 and casp-7. Homology between the active sites of casp-3 and casp-7 is extremely high with over 75% identity. The majority of the differences in active-site residues are localized to the non-prime binding pocket and these have been exploited in the design of small molecules selective for casp-3 over the other caspases using both canonical and unnatural amino acids ^24,25,36^. Such non-prime-side molecules have yet to be identified to selectively target casp-7. This barrier is primarily due to the superior activity of casp-3 over casp-7 against P4-P1 tetrapeptide-based substrates and inhibitors, as measured by *in vitro* kinetics assays. Importantly, these activity measurements do not include biologically relevant protein substrates, such as PARP, where casp-7 has improved degradation efficiency over casp-3 due to a unique casp-7 exosite for protein:protein interactions^47^.

Our characterization of non-hydrolyzable peptides revealed a potential and unexpected mechanism by which small molecules could selectivity bind casp-7 over casp-3. We exploited the subtle amino-acid substitution in the prime-side of the active site that includes targeting the Y151 side chain in casp-7 (*e*.*g*., F128 in casp-3) with ketomethylene inhibitors bearing a C-terminal PTAD. Our probe selectively labels casp-7 and will be a critical tool-like compound that will greatly assist with establishing the non-redundant roles of caspase family members.

We posit that future efforts will focus on optimization of the warhead chemistry for improved reactivity toward Y151 for the design of effective casp-7 inhibitors and not solely for probe-based detection. Understandably, this reactivity must be carefully tuned to avoid unwanted off-target promiscuity. In addition, we are developing shorter and more practical synthetic routes to generate dipeptidomimetic building blocks amendable to P1’ diversification. This methodology is applicable to the generation of inhibitors to structurally characterize the prime side of other proteases, as well as exploit the corresponding cysteine found at this position for the human inflammatory caspases-1, -4 and -5 (Figure 4C).

## Experimental Section

### Preparation of recombinant caspases

Casp-3, casp-7, and casp-8 were expressed and purified as previously described^48^.

### Enzymatic kinetic inhibition studies

Substrates, inhibitors and enzymes were diluted into an assay buffer containing 20 mM PIPES, pH 7.4, 10 mM NaCl, 1 mM EDTA, 10 mM DTT, and 10% sucrose. A 20 µL (2.5x) mixture of AMAC substrate (final concentration 100 µM; Ac-DEVD-AMAC for casp-3 and casp-7, and Ac-IETD-AMAC for casp-8 and inhibitor was dispensed into a black 96-well Costar flat-bottom polystyrene plate. 60 µL (1.67 x) enzyme solution (final concentration: 1 nM casp-3, 5 nM casp-7, and 25 nM casp-8) was added, mixed for 10 sec at 1000 rpm with fluorescence immediately read on a PerkinElmer EnVision plate reader with a λ_ex_ = [320] and λ_em_ = [460]. Data was analyzed using GraphPad Prism to determine k_inact_ and K_i_ in accordance with procedures previously outlined^49^. However, progress curves were fit 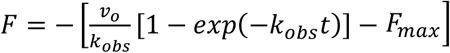 to account for the loss in fluorescence of AMAC substrates.

### Apoptotic protection assays

Jurkat A3 cells were cultured as described by ATCC using 10% FBS and pen/strep/glutamine at 37 °C with 5% CO_2_. Cells were plated into sterile Nunc Edge 96-well tissue culture-treated plates at a density of 10,000 cells/well and preincubated (or chased) with 20 µM of the indicated inhibitor or DMSO vehicle for the indicated time before (or after) induction with 10 ng/mL MegaFasL (AdipoGen® Life Sciences). 3 h after induction, cellular viability was measured using Cell-TiterGlo® on a PerkinElmer EnVision plate reader according to manufacturer’s protocol.

### X-ray crystallography of inhibitor-caspase complexes

Casp-3, casp-7 and casp-8 were incubated at 250-300 µM with a two-fold molar excess of Ac-DW3-KE, Ac-DEVD-Propionate, or Ac-DEVD-Propionate-AAA in 20 mM Tris, pH 8.0, 150 mM NaCl, 10 mM DTT, and 0.02% NaN_3_ for 2 h at 298 K and used immediately for co-crystallization experiments. Crystals of casp-8 in complex with Ac-DW3-KE were grown at 295 K after 1:1 dilution with 100 mM HEPES, 1.0 M sodium citrate, pH 8.1. Crystals of casp-3 in complex with Ac-DEVD-Propionate were grown at 295 K after 1:1 dilution with 0.10 M sodium citrate, pH 5.5, 10.0 % PEG6000, 10 mM DTT, 0.02% NaN_3_. Crystals of casp-3 in complex with Ac-DEVD-Propionate-AAA were grown at 295 K after 1:1 dilution with 0.10 M sodium citrate, pH 4.5, 12.0 % PEG6000, 10 mM DTT, 0.02% NaN_3_. Crystals of casp-7 in complex with Ac-DEVD-Propionate were grown at 295 K after 1:1 dilution with 0.15 M sodium citrate, 1.4 M sodium formate, pH 5.2. Crystals of casp-7 in complex with Ac-DEVD-Propionate-AAA were grown at 295 K after 1:1 dilution with 0.15 M sodium citrate, pH 5.6, 1.45 M sodium formate. His_6_-tags were not removed as proteins crystallized readily.

Data was collected on single, flash-cooled crystals at 100 K with a cryoprotectant consisting of mother liquor and 20% PEG400 (casp-3) or 25% glycerol (casp-7, casp-8) and were processed with HKL2000 in space groups P3_1_21 (casp-8), P2_1_2_1_2_1_ (casp-3:Ac-DEVD-Propionate), P2_1_2_1_2 (casp-3:Ac-DEVD-Propionate-AAA), and P322_1_ (casp-7)^50^. The asymmetric unit for casp-8:Ac-DW3-KE crystals contained one monomer of the biologically relevant homodimer, the complete biologically relevant homodimer for casp-3:Ac-DEVD-Propionate, casp-7:Ac-DEVD-Propionate, and caspase-7:Ac-DEVD-Propionate-AAA and one monomer per relevant homodimer for casp-3:Ac-DEVD-Propionate-AAA.

X-ray data was collected to 1.48 (casp-8:Ac-DW3-KE), 1.50 (casp-3:Ac-DEVD-Propionate), 2.60 (casp-7:Ac-DEVD-Propionate), 2.17 (casp-3:Ac-DEVD-Propionate-AAA)and 2.45 (casp-7:Ac-DEVD-Propionate-AAA) Å resolution on beamline 9.2 at the Stanford Synchrotron Radiation Lightsource (SSRL) (Menlo Park, CA). Data collection and processing statistics are summarized in Tables S1-S3.

### Structure solution and refinement

All caspase structures were determined by molecular replacement (MR) with Phaser^51,52^ using previously published casp-3 (4JJE), casp-7 (4JJ8), and casp-8 (4JJ7) as the initial respective search models. All structures were manually built with Coot^53^ and iteratively refined using Phenix^54^ with cycles of conventional positional refinement. For the low-resolution casp-7 structures, non-crystallography symmetry (NCS) restraints between the two subunits of the homodimer were applied during refinement. For all structures, the electron density maps clearly identified the peptide ligands and covalent attachment to the caspase active-site cysteine. Water molecules were automatically positioned by Phenix using a 2.5σ cutoff in *f*_o_-*f*_c_ maps and manually inspected.

The final R_cryst_ and R_free_ are 14.9% and 16.7%, respectively, for casp-8:Ac-DW3-KE, 17.2% and 19.7% for casp-3:Ac-DEVD-Propionate, 18.3% and 21.1% for casp-7:Ac-DEVD-Propionate, 22.0% and 26.3% for casp-3:Ac-DEVD-Propionate-AAA, and 17.9% and 21.5% for casp-7:Ac-DEVD-Propionate-AAA. All models were validated with the PDB server prior to deposition and all residues for the structures are in the most favorable and allowed regions in the Ramachandran plot. Coordinates and structure factors have been deposited in the Protein Data Bank with accession entries 6X8H (casp-8:Ac-DW3-KE), 6X8I (casp-3:Ac-DEVD-Propionate), 6X8J (casp-7:Ac-DEVD-Propionate), 6X8K (casp-3:Ac-DEVD-Propionate-AAA), and 6X8L (casp-7:Ac-DEVD-Propionate-AAA). Structure refinement statistics are provided in Tables S1-S3.

### Labeling of caspase-3 and -7 with biotin-DEVD-Propionate-3/4-PTAD

Prechilled biotin-DEVD-Propionate-4-PTAD or biotin-DEVD-Propionate-3-PTAD (10 µL, 10 mM) in DMF was treated with prechilled dibromantin (10 µL, 10 mM) in DMF to generate corresponding triazole active species and immediately used to label 250 nM caspase-3 or caspase-7 in caspase activity buffer (50 mM HEPES pH 7.4, 0.1% CHAPS, 10 mM KCl, 50 mM sucrose, 1 mM MgCl_2_, 10 mM DTT) at the indicated inhibitor concentrations. The samples were incubated overnight at room temperature and then proteins were resolved by SDS-PAGE, transferred to PVDF membranes, blocked with 5% non-fat dry milk in TBST (0.1% Tween-20), probed with IRDye® 800CW Streptavidin (LI-COR 926-32230, 1:10,000, 4 °C, overnight), washed (rt, 3 × 5 min, TBST) and visualized on a LICOR Odyssey Scanner.

### Synthetic Protocols

Semicarbazide resin and AMAC-based substrates were synthesized as previously^55-57^. Detailed protocols for the synthesis of all molecules and inhibitors are outlined in the Supporting Information.

## Supporting information

Solania et al 2024 Supporting Information

## Associated Content

### Supporting information

Supporting information, including co-complex caspase crystal structures and synthesis and characterization of the peptidomimetics presented are available. Structures of casp-8:Ac-DW3 (PDB ID 6X8H), casp-3:Ac-DEVD-Propionate (PDB ID 6X8I), casp-7:Ac-DEVD-Propionate (PDB ID 6X8J), casp-3:Ac-DEVD-Propionate-AAA (PDB ID 6X8K), and casp-7:Ac-DEVD-Propionate-AAA (PDB ID 6X8L) have been deposited in the Protein Database (PDB).

### Author contributions

A.S. conceived the study. D.W.W. supervised the project. A.S. prepared all materials and performed all assays. A.S. and J.H.X. collected the x-ray crystallographic data and determined the co-crystal structures. The manuscript was written by A.S. and D.W.W. with contributions from J.H.X. All authors have given approval to the final version of the manuscript.

### Funding Sources

The authors gratefully acknowledge financial support from The Scripps Research Institute and the U.S. National Institutes of Health 5R21AI112796, 1R01GM118382, and 1R35GM136286 (to D.W.W.).

### Notes

The authors declare no competing financial interests.

## Acknowledgement

We thank I. Wilson, R. Stanfield, M. Elsliger, and X. Dai for x-ray data collection and computational assistance; H. Rosen and L. Lairson for access to instrumentation; and the staff of the Stanford Synchrotron Radiation Lightsource.

## Abbreviations

AEBSF: aminoethylbenzylsulfonic acid
AOMK: acyloxymethylketone(s)
casp: caspase
CMK: chloromethylketone
DFP: diisopropylfluorophosphate
FMK: fluoromethylketone
KE: thiophene carboxylic acid
PTAD: 4-phenyl-1,2,4-triazole-3,5-dione
SPPS: solid-phase peptide synthesis
TS: transition-state

---

**Scheme 1.**
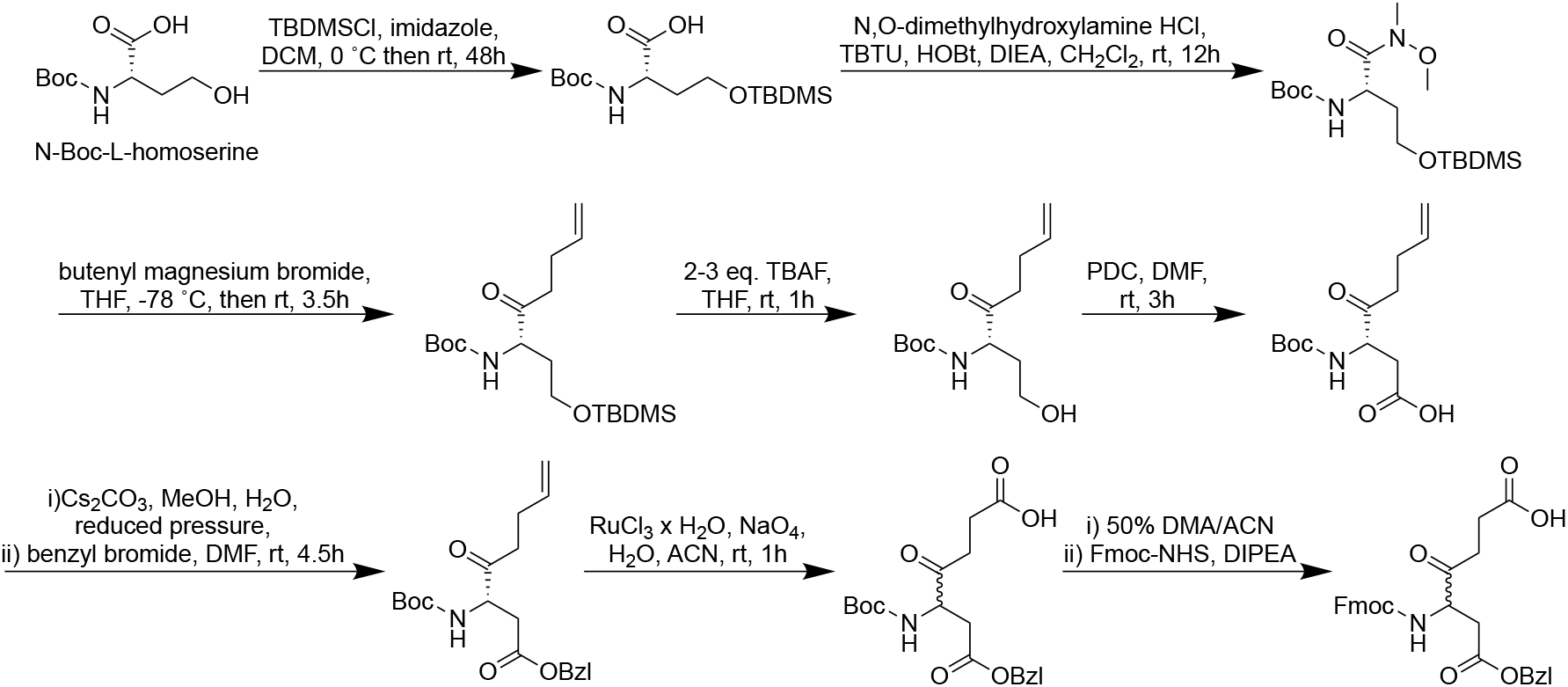
Synthesis of Fmoc-Asp(OBzl)-Propionate.

**Scheme 2.**
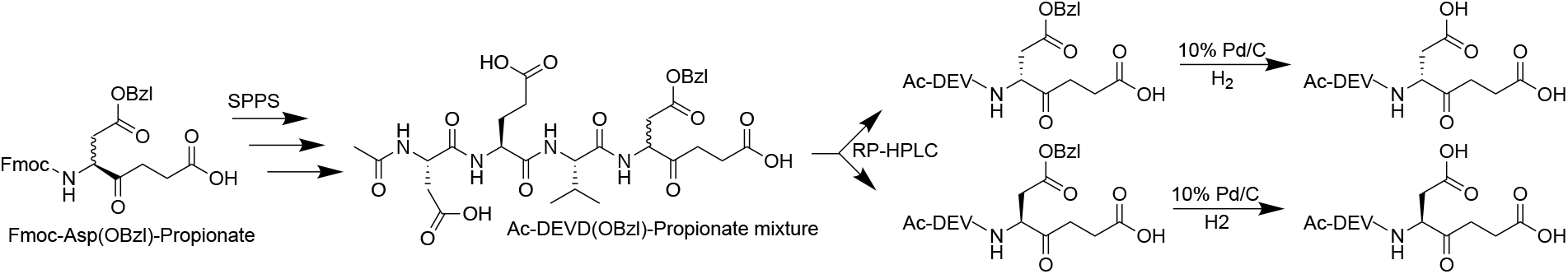
Synthesis and purification of Ac-DEVD-Propionate isomers.

**Scheme 3.**
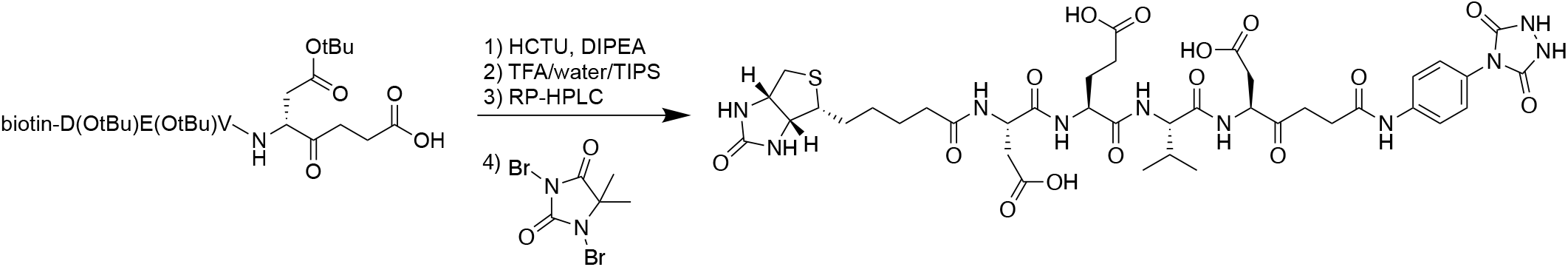
Synthesis of biotin-DEVD-Propionate-4-PTAD.

