## Supplementary material for "Caspase prime-side active-site characterization with non-hydrolyzable peptides assists in the design of a caspase-7-selective irreversible probe": Solania et al 2024 Supporting Information

**Table of Contents**

S2-S8 Detailed synthetic methods and compound characterization

S9 Figure S1: Structure of casp-8 in complex with Ac-DW3-KE

S10 Figure S2: Dose responses of Ac-IETD-Propionate against casp-3, casp-7, and casp-8

S11 Figure S3: Structures of casp-3 and casp-7 in complex with Ac-DEVD-Propionate

S12 Figure S4: Structures of casp-3 and -7 in complex with Ac-DEVD-Propionate-AAA

S13 LC-MS traces of synthesized peptides Ac-DEV(*L/D*)D-Propionate

S14 Table S1: Structure statistics for casp-8:Ac-DW3-KE complex

S15 Table S2: Statistics for structures of casp-3 and casp-7 in complex with Ac-DEVD-Propionate

S16 Table S3: Statistics for structures of casp-3 and casp-7 in complex with Ac-DEVD-Propionate-AAA

S17 Supporting information references

***Detailed synthetic methods and compound characterization***

Unless otherwise noted, all reagents and solvents were purchased from commercial suppliers and were used without further purification. All solution-phase reactions were performed in an inert atmosphere of dry nitrogen or argon. Silica gel column chromatography was performed using 60 Å silica gel (230−400 mesh). Reactions were monitored on TLC plates (silica gel 60, F254 coating, EMD Millipore, 1057150001), and spots were either monitored under UV light (254 mm) or stained with ninhydrin. NMR spectra recorded in $CDCl_3$ or $MeOD_4$ used TMS as the internal reference. When noted, compounds were purified via a Waters Preparative HPLC system using a 19x100mm or 19x150mm C18 5 µm OBD column.

**Boc-hSer(OTBDMS)-OH**

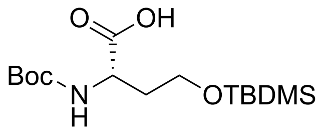


A flame dried 100 mL round bottom flask and stir bar was charged with Boc-hSer-OH (4.00 g, 18.25 mmol) and suspended in 60 mL anhydrous THF. The solution was chilled before TBDMSCl (5.50 g, 36.5 mmol) and imidazole (3.11 g, 46.61 mmol) were added. The reaction was stirred for 48 hr at room temperature (rt) under $N_2$. The solvent was evaporated, and the crude material resuspended in 40 mL MeOH. MeOH was evaporated and the residue resuspended with 200 mL DCM and the organic layer was washed with 0.1 M HCl (2 x 40 mL) and saturated brine. The organic layer was dried over magnesium sulfate, filtered and concentrated to yield Boc-hSer(OTBDMS)-OH (90%) as an oil, which was used without further purification.

**Boc-hSer(OTBDMS)-Weinrab**

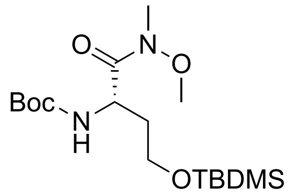


A flame dried 50 mL round bottom flask and stir bar was charged with Boc-hSer(OTBDMS)-OH (5.45 g, 16.34 mmol), PyBOP (8.5 g, 16.34 mmol) in 20 mL anhydrous DCM. The solution was chilled (0 °C) and DIPEA (4.27 mL, 3.17 g, 24.51 mmol) was added. In a separate flask, a solution of DMHA (1.99 g, 20.43 mmol) and DIPEA (4.27 mL, 3.17 g, 24.51 mmol) was sonicated until complete dissolution. The DMHA solution was then added dropwise to the chilled stirring solution of activated Boc-hSer(OTBDMS)-OH and allowed to react for 2 hr at rt under $N_2$. The solvent was evaporated, and the residue resuspended in 200 mL diethyl ether. The organic layer was washed with saturated bicarbonate and saturated brine. The organic layer was dried over magnesium sulfate, filtered and concentrated before purification on silica gel (2:1 hexanes: ethyl acetate) to provide Boc-hSer(OTBDMS)-Weinrab (71%) as an oil.

**TLC** (hexanes: ethyl acetate, 2:1 v/v): $R_f$ = 0.35.
**$^1$H NMR** (600 MHz, Chloroform-*d*) δ 5.50 (d, *J* = 9.3 Hz, 1H), 4.74 (d, *J* = 10.0 Hz, 1H), 3.78 (s, 3H), 3.74 – 3.59 (m, 2H), 3.20 (s, 3H), 1.96 (ddt, *J* = 14.2, 9.6, 4.8 Hz, 1H), 1.76 – 1.66 (m, 1H), 1.42 (s, 9H), 0.90 (q, *J* = 2.2 Hz, 9H), 0.05 (d, *J* = 7.4 Hz, 6H).
**LRMS (*m/z*):** $[M]^+$ calcd. for $C_{17}H_{36}N_2O_5Si$, 376.24; found, $[MH]^+$ 377.40.

**Boc-hSer(OTBDMS)-butene**

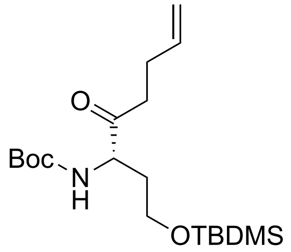


A flame dried 50 mL round bottom flask and stir bar was charged with magnesium pellets (2.58 g, 106 mmol) and purged with argon. 115 mL anhydrous THF was added and the suspension was chilled (0 °C) before the addition of 4-bromo-1-butene (7.55 mL, 10.0 g, 74.4 mmol) in portions and reacted at rt for 30 min. Under argon, the Grignard was added to a stirring solution of chilled (-78 °C) Boc-hSer(OTBDMS)-Weinrab (4.00 g, 10.62 mmol) in 150 mL anhydrous THF. After addition, the acetone:dry ice bath was removed, and the reaction was stirred at rt for 4 hr under argon. Approximately half of the THF was evaporated and the solution was chilled (0 °C) before being quenched dropwise by 5% ammonium chloride. After 5 mL, the cold bath was removed and 5% ammonium chloride (100 mL) was added. The aqueous layer was extracted with ethyl acetate (400 mL) and the organic layer was washed with saturated brine (100 mL), was dried over magnesium sulfate, filtered and concentrated before purification on silica gel (3:1 hexanes: ethyl acetate) to provide Boc-hSer(OTBDMS)-butene (92%) as an oil.

**TLC** (hexanes: ethyl acetate, 2:1 v/v): $R_f$ = 0.66.
**$^1$H NMR** (600 MHz, Chloroform-*d*) δ 5.84 – 5.71 (m, 2H), 5.00 (dq, *J* = 17.2, 1.7 Hz, 1H), 4.95 (dt, *J* = 10.2, 1.5 Hz, 1H), 4.30 – 4.24 (m, 1H), 3.66 (t, *J* = 5.6 Hz, 2H), 2.66 (dt, *J* = 17.6, 7.5 Hz, 1H), 2.59 (dt, *J* = 17.6, 7.4 Hz, 1H), 2.31 (tdd, *J* = 7.8, 6.6, 1.4 Hz, 2H), 2.00 (dq, *J* = 15.6, 5.4, 4.7 Hz, 1H), 1.91 – 1.77 (m, 1H), 1.41 (s, 9H), 0.87 (s, 9H), 0.02 (d, *J* = 3.8 Hz, 6H).
**$^{13}$C NMR** (151 MHz, Chloroform-*d*) δ 208.88, 155.61, 137.04, 115.28, 79.49, 70.92, 59.86, 58.29, 38.49, 36.51, 33.32, 28.33, 28.31, 27.46, 25.86, -5.66.
**LRMS (*m/z*)**: $[M]^+$ calcd. for $C_{19}H_{37}NO_4Si$, 371.25; found, $[MH]^+$ 372.34.

**Boc-Asp(OBzl)-butene**

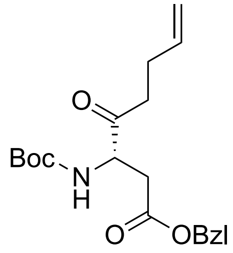


*Step 1:* Boc-hSer(OTBDMS)-butene (3.65 g, 9.82 mmol) was dissolved in 25 mL THF and tetrabutylammonium fluoride (TBAF, 1M in THF, 14.8 mL, 14.8 mmol) was added in one batch to the stirring solution of amino acid and stirred at rt for 1 h. The solvent was evaporated and resuspended with diethyl ether. The organic layer and washed with saturated $NH_4Cl$ (30 mL) and saturated brine. The organic layer was dried over magnesium sulfate, filtered and concentrated to yield Boc-hSer(OH)-butene in quantitative yield as an oil, which was used without further purification.

*Step 2:* Boc-hSer(OH)-butene (2.53 g, 9.82 mmol) was dissolved in 18 mL freshly distilled DMF and pyridinium dichromate (11.09 g, 29.47 mmol) was added in 1 batch. This solution was stirred at rt for 2.5 hr before quenching with 100 mL saturated $NaHCO_3$. The aqueous layer was washed with hexanes (2 x 60 mL), acidified with conc. HCl to pH 2, and extracted diethyl ether (2 x 100 mL). The combined organic layer was concentrated and used without further purification.

*Step 3:* Boc-Asp(OH)-butene in 20 mL solution of 9:1 MeOH:$H_2O$ was added 20% $CsCO_3$ until pH reached 8 and then the solvent evaporated to near dryness. The residue was dissolved in fresh anhydrous DMF (18 mL) and evaporated to dryness (2x). The dried residue was dissolved in 18 mL fresh anhydrous DMF and benzyl bromide (1.34 mL, 1.93 g, 11.30 mmol) was added. The solution was reacted at rt for 6 h. The solvent was evaporated, and the residue resuspended with ethyl acetate (200 mL) and washed with saturated brine (3 x 75 mL). The combined organic layer was dried over magnesium sulfate, filtered and concentrated. The crude product was purified on silica gel (7:1 to 4:1 hexanes: ethyl acetate) to provide Boc-Asp(OBzl)-butene in 49 % yield (3 steps).

**TLC** (Hexanes: ethyl acetate, 4:1 v/v): $R_f$ = 0.40.
**$^1$H NMR** (600 MHz, Chloroform-*d*) δ 7.40 – 7.30 (m, 5H), 5.78 (ddt, *J* = 16.8, 10.2, 6.5 Hz, 1H), 5.59 (d, *J* = 9.0 Hz, 1H), 5.15 – 5.06 (m, 2H), 5.02 (dq, *J* = 17.1, 1.7 Hz, 1H), 4.97 (dt, *J* = 10.3, 1.5 Hz, 1H), 4.45 (dt, *J* = 9.3, 4.8 Hz, 1H), 3.03 (dd, *J* = 17.1, 4.7 Hz, 1H), 2.80 (dd, *J* = 17.0, 5.0 Hz, 1H), 2.74 – 2.59 (m, 2H), 2.31 (qdd, *J* = 6.7, 2.9, 1.6 Hz, 2H), 1.45 (s, 9H).
**$^{13}$C NMR** (151 MHz, Chloroform-*d*) δ 207.86, 171.47, 155.51, 136.97, 135.42, 128.73, 128.54, 128.37, 115.45, 80.42, 66.98, 55.98, 38.24, 35.89, 28.42, 27.47.
**LRMS (*m/z*):** $[M]^+$ calcd. for $C_{20}H_{27}NO_5$, 361.19; found, $[MH]^+$ 362.29.

**Fmoc-Asp(OBzl)-Propionate**

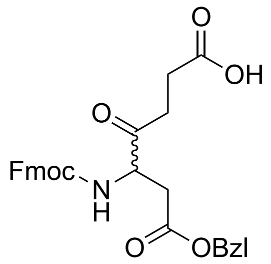


*Step 1:* Boc-Asp(OBzl)-butene (1.75 g, 4.86 mmol) in 50 mL acetonitrile was added a solution of ruthenium (III) chloride hydrate (0.030 g, 0.15 mmol) and sodium periodate (7.27 g, 34.0 mmol) in 10 mL $H_2O$. This was reacted at rt for 1 hr before the pH was adjusted to pH ~2 with 1 M HCl. The aqueous layer was diluted with saturated brine (50 mL) and extracted with ethyl acetate (2 x 100 mL). The organic layer was washed with saturated brine and dried over magnesium sulfate, filtered, and dried thoroughly to yield Boc-Asp(OBzl)-Propionate, which was used directly for the next step.

*Step 2:* Boc-Asp(OBzl)-Propionate (1.48 g, 3.90 mmol) was suspended in 33% TFA in DCM until and the reaction monitored by TLC (10% MeOH in DCM). The solvent was evaporated and triturated with another aliquot of DCM (2x 25 mL) to yield $NH_2$-Asp(OBzl)-Propionate, which was used directly in the next step.

*Step 3:* A solution of $NH_2$-Asp(OBzl)-Propionate in 10 mL anhydrous DCM was chilled (0 °C) and was added Fmoc-NHS (1.38 g, 3.90 mmol) and triethylamine (1.36 mL, 0.99 g, 9.76 mmol). After addition, the reaction was stirred at rt for 3 h. The solvent was evaporated, and the residue resuspended in diethyl ether (50 mL). The organic layer was washed with 1 M HCl (100 mL). The aqueous layer was extracted with another aliquot of diethyl ether (50 mL) and the combined organic layer was washed with saturated brine, dried over magnesium sulfate, filtered and concentrated. The crude product was purified on silica gel (4 to 8% MeOH in DCM gradient) to afford Fmoc-Asp(OBzl)-Propionate (0.98 g, 50% yield, 3 steps).

**TLC** (dichloromethane: methanol, 9:1 v/v): $R_f$ = 0.40.
**LRMS (*m/z*):** $[M]^+$ calcd. for $C_{29}H_{27}NO_7$, 501.18; found, $[MH]^+$ 502.31.

**Ac-DEV(*L*)D(OBzl)-Propionate and Ac-DEV(*D*)D(OBzl)-Propionate**

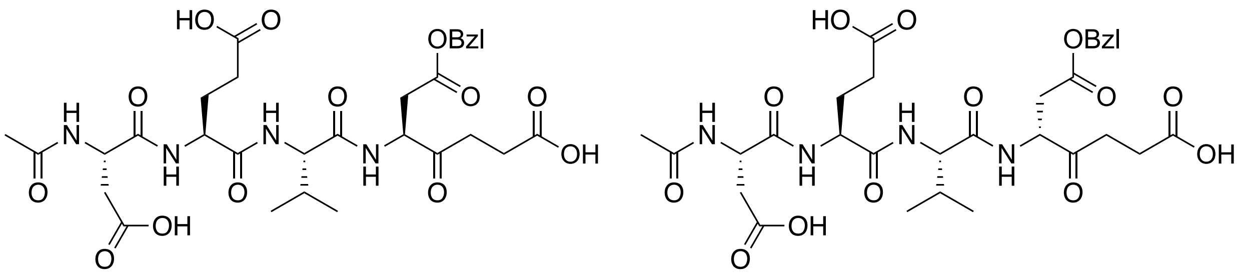

2-chlorotrityl chloride resin (1.0 g, 1.0-1.4 mmol/g) was pre-swelled for 30 min with 10 mL anhydrous DCM. This solution was discarded and Fmoc-Asp(OBzl)-Propionate (200 mg, 0.40 mmol) and DIPEA (700 µL, 517 mg, 4 mmol) in 10 mL anhydrous DCM was added, and the resin was nutated for 90 min at rt. The loading solution was discarded, and the resin washed twice with DCM and DMF. This was performed in between all subsequent steps. Deprotection was performed with 20% pyrrolidine (20 min). Coupling was performed using N-Fmoc-protected amino acids, HCTU, and DIPEA at a 3:3:10 ratio with respect to the loading of the resin, which was determined by Fmoc quantitation. The deprotection, coupling, and washing steps were repeated with the corresponding amino acids until the desired length was achieved. Prior to cleavage, the resin was additionally washed with DCM and MeOH 3x before being dried *in vacuo.* Cleavage was performed using 1 mL/100 mg resin of TFA/water/TIPS at a 90:5:5 ratio for 1 hr. The resin was washed with another aliquot of cleavage cocktail and the combined cleavage solutions were concentrated before precipitation with cold diethyl ether. The pellet was dried under a stream of $N_2$ and dissolved a minimal volume of DMSO before purification by preparatory reverse phase HPLC (19x150 mm XBridge C18, $CH_3CN/H_2O$/0.1% TFA, 25:75 to 90:10 over 13 min; 20 mL/min) and lyophilization to yield Ac-DEV(*L*)D(OBzl)-Propionate and Ac-DEV(*D*)D(OBzl)-Propionate (19.1 mg and 17.9 mg, respectively).

**LRMS (*m/z*):** $[M]^+$ calcd. for $C_{30}H_{40}N_4O_{13}$, 664.26; found, $[MH]^+$ 665.25 was found for both fractions 11 and 12 (see LC-MS trace below).

**Ac-DEV(*L*)D-Propionate (Ac-DEVD-Propionate) and Ac-DEV(*D*)D-Propionate**

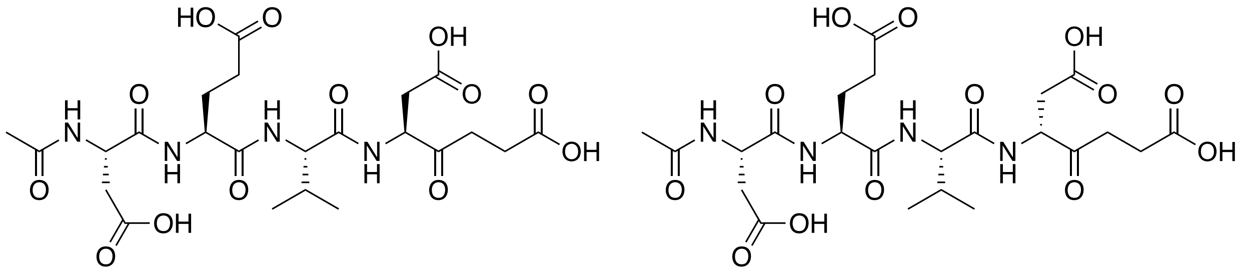

Ac-DEV(*L*)D(OBzl)-Propionate and Ac-DEV(*D*)D(OBzl)-Propionate was individually dissolved in 5 mL MeOH and treated with 10% Pd/C (5 mg) in 1 mL toluene. The vials were purged with argon (2 x) and $H_2$ (2 x) and reacted at rt for 4 hr under $H_2$ balloon. The Pd/C was removed using prewetted Celite® and the solvent evaporated. The residue was resuspended in 10% acetonitrile in water and purified by reversed phase HPLC (19x150 mm XBridge C18, $CH_3CN/H_2O$/0.1% TFA, 100:0 to 50:50 over 13 min; 20 mL/min) and lyophilization to yield Ac-DEV(*L*)D-Propionate (**Ac-DEVD-Propionate**) and Ac-DEV(*D*)D-Propionate (12.4 mg and 11.3 mg, respectively).

**LRMS (*m/z*):** $[M]^+$ calcd. for $C_{23}H_{34}N_4O_{13}$, 574.21; found, $[MH]^+$ 575.10 was found for both products.

**Ac-IET(*L/D*)D-Propionate (racemic mixture)**

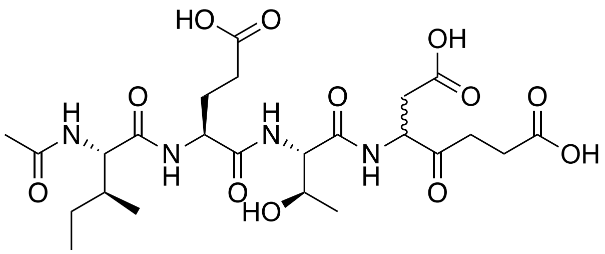

Ac-IETD-Propionate was synthesized similarly to Ac-DEVD-Propionate; however, Ac-IET(*D/L*)D(OBzl)-Propionate diastereomers could not be readily isolated and were used as a racemic mixture for the *in vitro* enzymatic analyses against casp-3, -7, and -8. Resolution of the two diastereomers was not pursued due to poor inhibition (Fig. S2).

**LRMS (*m/z*):** $[MH]^+$ calcd. for $C_{24}H_{38}N_4O_{12}$, 575.26; found, 575.07.

**Ac-DEVD-Propionate-AAA**

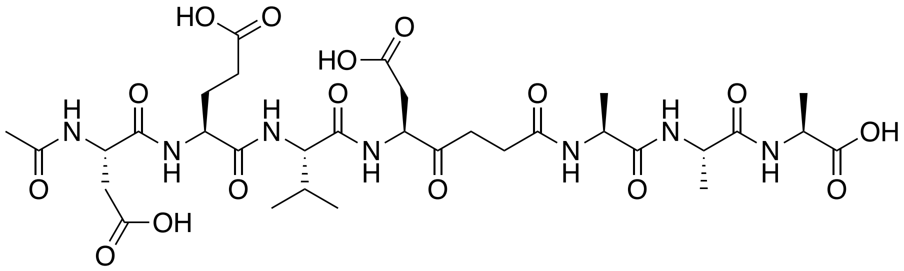

Ac-DEVD-Propionate-AAA was prepared similarly to Ac-DEVD-Propionate with the except Fmoc-Ala-OH was loaded onto 2-chlorotrityl chloride resin and Fmoc-Asp(OtBu)-Propionate was used in place of Fmoc-Asp(OBzl)-Propionate.

**LRMS (*m/z*):** $[M]^+$ calcd. for $C_{32}H_{49}N_7O_{16}$, 787.32; found, $[MH]^+$ 788.32.

**4-(4-aminophenyl)-1,2,4-triazolidine-3,5-dione (4NH$_2$-PTAD)**

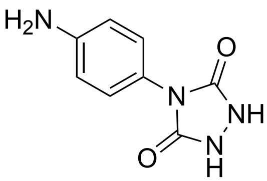

*Step 1:* A 100 mL round bottom was charged with 4-nitrophenylisocyanate (1000 mg, 6.093 mmol) in 15 ml anhydrous toluene and chilled (0 °C, 15 min). Meanwhile, ethyl carbazate (634 mg, 6.093 mmol) was dissolved in 15 mL anhydrous toluene in a 20 mL scintillation vial. The ethyl carbazate was chilled (0 °C, 15 min) before dropwise adding to stirring isocyanate. The ice bath was removed, and the reaction stirred at rt for 2 hr. The stirring solution was then chilled (0 °C, 15 min) and the pale-yellow powder collected by vacuum filtration. Residual toluene was removed to yield ethyl 2-((4-nitrophenyl)carbamoyl)hydrazine-1-carboxylate and used directly for the next step.

*Step 2:* A 20 mL scintillation vial charged with ethyl 2-((4-nitrophenyl)carbamoyl)hydrazine-1-carboxylate dissolved in 4 M potassium hydroxide (923 mg KOH in 4.1 mL water, 16.5 mmol) was refluxed (100 °C, 4 hr). The solution was allowed to cool to the touch and filtered while warm. The filtrate was allowed to cool to rt and then acidified with 1 N $HCl_{(aq.)}$ to pH 1. The product was then collected by vacuum filtrated to yield

4-nitrophenyl-1,2,4-triazolidine-3,5-dione (55% over 2 steps) as a solid white powder and used without further purification.

*Step 3:* A 50 mL round bottom flask charged with 4-(4-nitrophenyl)-1,2,4-triazolidine-3,5-dione (740 mg, 3.33 mmol) was dissolved in 14 mL MeOH. The flask was sealed with an inverted septum, purged of air, then backfilled with $H_2$ via balloon. This cycle was repeated, and the reaction vigorously stirred under $H_2$ for 3 hr at room temperature. The solution was filtered through a bed of prewetted Celite to remove Pd/C. The solvent was removed *in vacuo* before iterative silica gel column chromatography (20% MeOH in DCM) to yield 4-(4-aminophenyl)-1,2,4-triazolidine-3,5-dione (78%) as a brown powder.

**$^1$H NMR** (500 MHz, Methanol-$d_4$) δ 7.11 – 7.05 (m, 2H), 6.79 – 6.74 (m, 2H).
**LRMS (*m/z*):** $[M]^+$ calcd. for $C_8H_8N_4O_2$, 192.06; found, $[MH]^+$ 193.28.

**4-(3-aminophenyl)-1,2,4-triazolidine-3,5-dione (3NH$_2$-PTAD)**

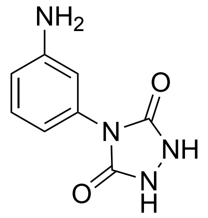


4-(3-aminophenyl)-1,2,4-triazolidine-3,5-dione was synthesized similarly to 4-(4-aminophenyl)-1,2,4-triazolidine-3,5-dione starting from 3-nitrophenylisocyanate in 35% overall yield (3 steps) as a pale tan powder.

**$^1$H NMR** (500 MHz, Methanol-$d_4$) δ 7.22 – 7.14 (m, 1H), 6.73 (dd, $J$ = 7.9, 1.2 Hz, 2H), 6.71 – 6.63 (m, 1H).
**LRMS (*m/z*):** $[M]^+$ calcd. for $C_8H_8N_4O_2$, 192.06; found, $[MH]^+$ 193.27.

**Biotin-D(OtBu)E(OtBu)VD(OtBu)-Propionate**

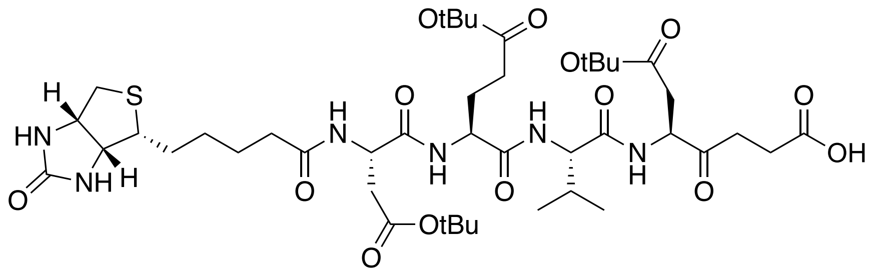


Biotin-D(OtBu)E(OtBu)VD(OtBu)-Propionate was prepared similarly to Ac-DEVD-Propionate-AAA, except cleavage was performed to yield the full protected peptide using a 1:1:8 solution of acetic acid:trifluoroethanol:DCM (2 × 8 mL, 30 min each). The cleavage solutions were combined and diluted with 10 volumes hexane. Acetic acid was removed via azeotropic evaporation. The residue was resuspended in neat DMSO and purified by reversed phase HPLC (19x150 mm XBridge C18, $CH_3CN/H_2O$/0.1% TFA, 65:35 to 0:100 over 13 min; 20 mL/min) and lyophilization to yield biotin-D(OtBu)E(OtBu)VD(OtBu)-Propionate as a fluffy white pellet.

**LRMS (*m/z*):** $[M]^+$ calcd. for $C_{43}H_{70}N_6O_{14}S$, 926.47; found, $[MH]^+$ 927.64.

**Biotin-DEVD-Propionate-4-PTAD**

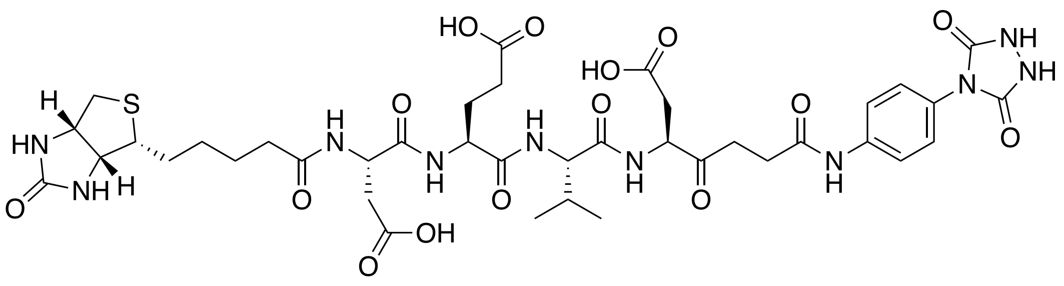

A 50 mL conical was charged with biotin-D(OtBu)E(OtBu)VD(OtBu)-Propionate (20 mg, 0.021mmol) in 862 µL DMSO, HCTU (35.7 mg, 0.86 mmol), DIPEA (16.7 mg, 0.13 mmol) and stirred for 5 min at rt. $4NH_2$-PTAD (13.7 mg, 0.24 mmol) was then added as a solid to the stirring solution. After 1 hr at rt, the peptide was treated with a solution of 90:5:5 trifluoroacetic acid/water/triisopropylsilane (7 mL) for 1 hr at rt. The solution was then concentrated to ~4 mL with gentle stream of $N_2$ and diluted up to 30 mL with diethyl ether. The suspension was vortexed and iced for 30 min. The product was centrifuged (4,000 x g, 5 min). The supernatant was discarded and the pellet resuspended in a minimal amount of 50% acetonitrile in water before dilution with water to a final concentration of 25% acetonitrile before purification by reversed phase HPLC (19x150 mm XBridge C18, $CH_3CN/H_2O$/0.1% TFA, 90:10 to 50:50 over 21 min; 25 mL/min) and lyophilization to yield biotin-DEVD-Propionate-4-PTAD as a fluffy white pellet (4.3 mg).

**LRMS (*m/z*):** $[M]^+$ calcd. for $C_{39}H_{52}N_{10}O_{15}S$, 932.33; found, $[MH]^+$ 933.74.

**Biotin-DEVD-Propionate-3-PTAD**

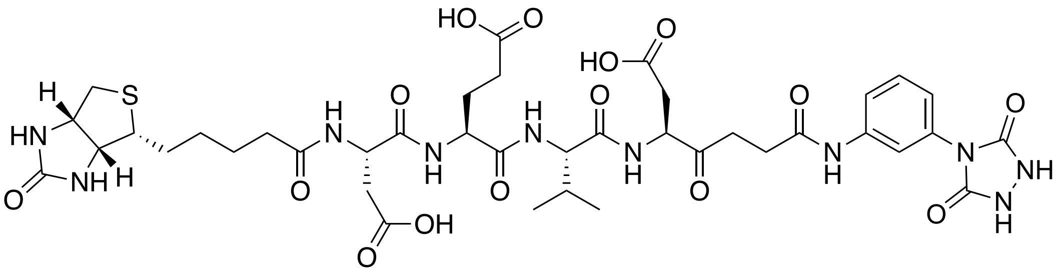

Biotin-DEVD-Propionate-3-PTAD was synthesized similarly to Biotin-DEVD-Propionate-3-PTAD with the exception that $3NH_2$-PTAD was used in place of $4NH_2$-PTAD (9.2 mg).

**LRMS (*m/z*):** $[M]^+$ calcd. for $C_{39}H_{52}N_{10}O_{15}S$, 932.33; found, $[MH]^+$ 933.73.

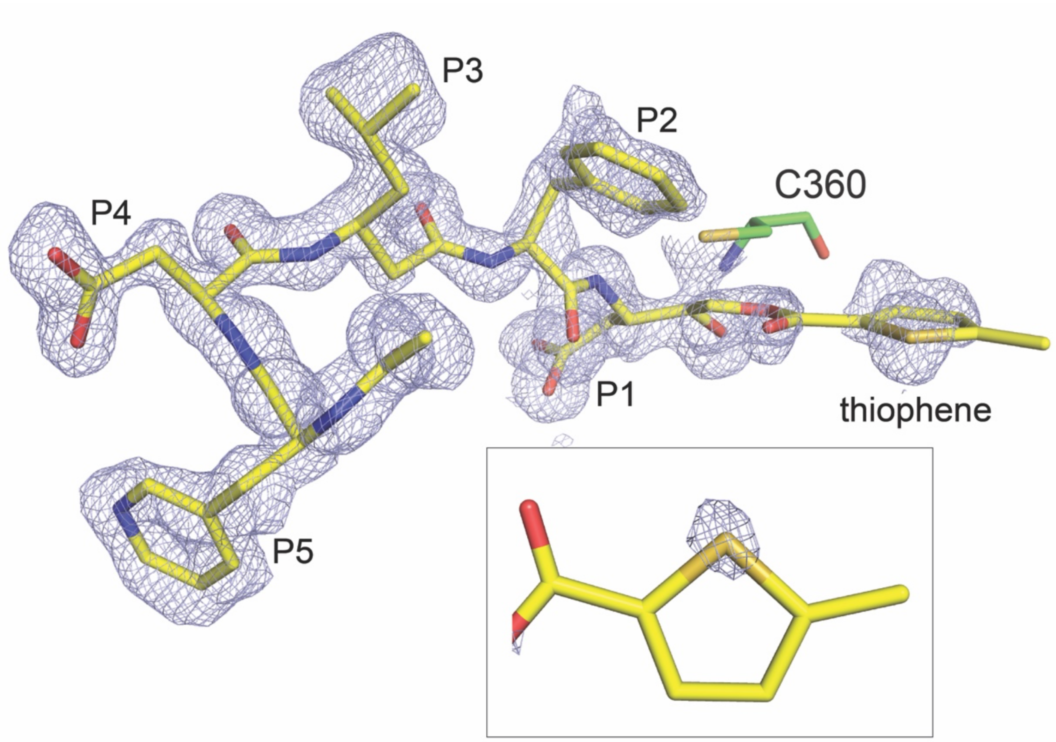


**Figure S1**. Casp-8 in complex with Ac-DW3-KE. (A) Naïve $f_o$-$f_c$ density (blue) of Ac-DW3-KE (yellow carbon) bound to casp-8 (green carbon) contoured at 1σ shows that the 5-methyl-2-thiophene carboxylate leaving group remains bound and occupies the S1' subsite (blue nitrogen, red oxygen, mustard sulfur). (Inset) Naïve $f_o$-$f_c$ density (blue) of Ac-DW3-KE thiophene contoured at 2.5σ depicting a strong electron density corresponds with placement of the thiophene sulfur atom.

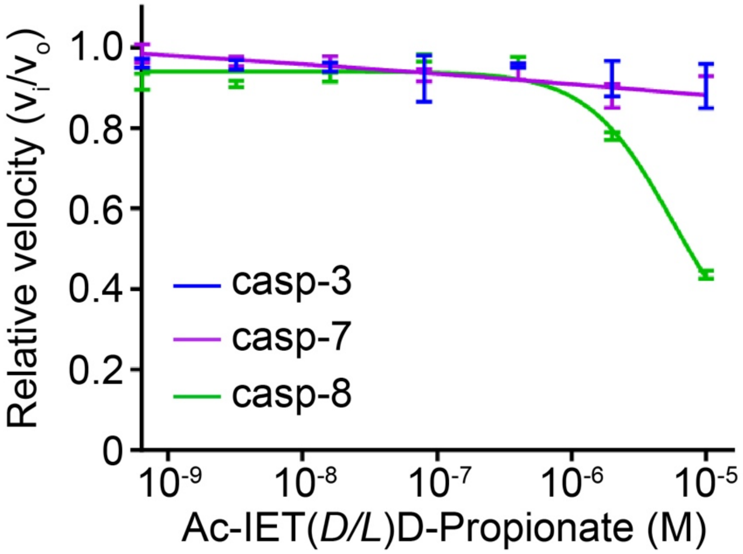


**Figure S2.** Activity dose response curves of Ac-IET(*D/L*)D-Propionate against panel of apoptotic caspases shows that replacement of the DEVD tetrapeptide sequence with the canonical recognition sequence IETD for casp-8 did not improve inhibition efficiency against casp-8. The change in peptide sequence completely ablated inhibition against casp-3 and -7.

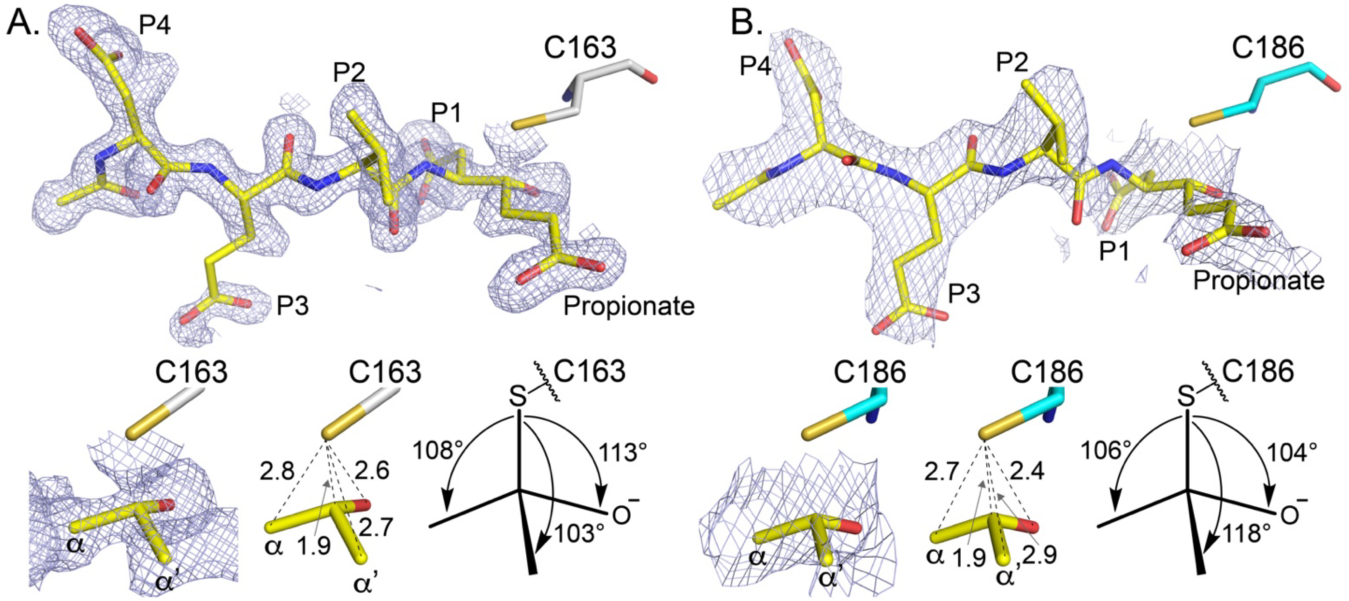


**Figure S3.** Co-crystal structures of casp-3 and casp-7 in complex with Ac-DEVD-Propionate determined to 1.50 Å and 2.60 Å, respectively. (A) Naïve $f_o$-$f_c$ density (blue) of Ac-DEVD-Propionate (yellow carbon) contoured at 1σ bound to casp-3 (grey carbon) shows highly resolvable density for the propionate group within the caspase S1’ subsite (blue nitrogen, red oxygen, mustard sulfur). Bottom panels include: zoomed in view of naïve density of casp-3:Ac-DEVD-Propionate depicting the *sp3*-tetrahedral adduct; distances between the casp-3 C163 sulfur and carbons of Ac-DEVD-propionate; and the angles formed between casp-3 C163 sulfur and three other substituents of P1 carbon 1 that fall within range of the ideal 109.5° for tetrahedral geometry. (B) Naïve $f_o$-$f_c$ density (blue) of Ac-DEVD-Propionate (yellow carbon) contoured at 1σ bound to casp-7 (cyan carbon) demonstrating strong density for propionate in the S1’ subsite. Bottom panels are depicted as in (A).

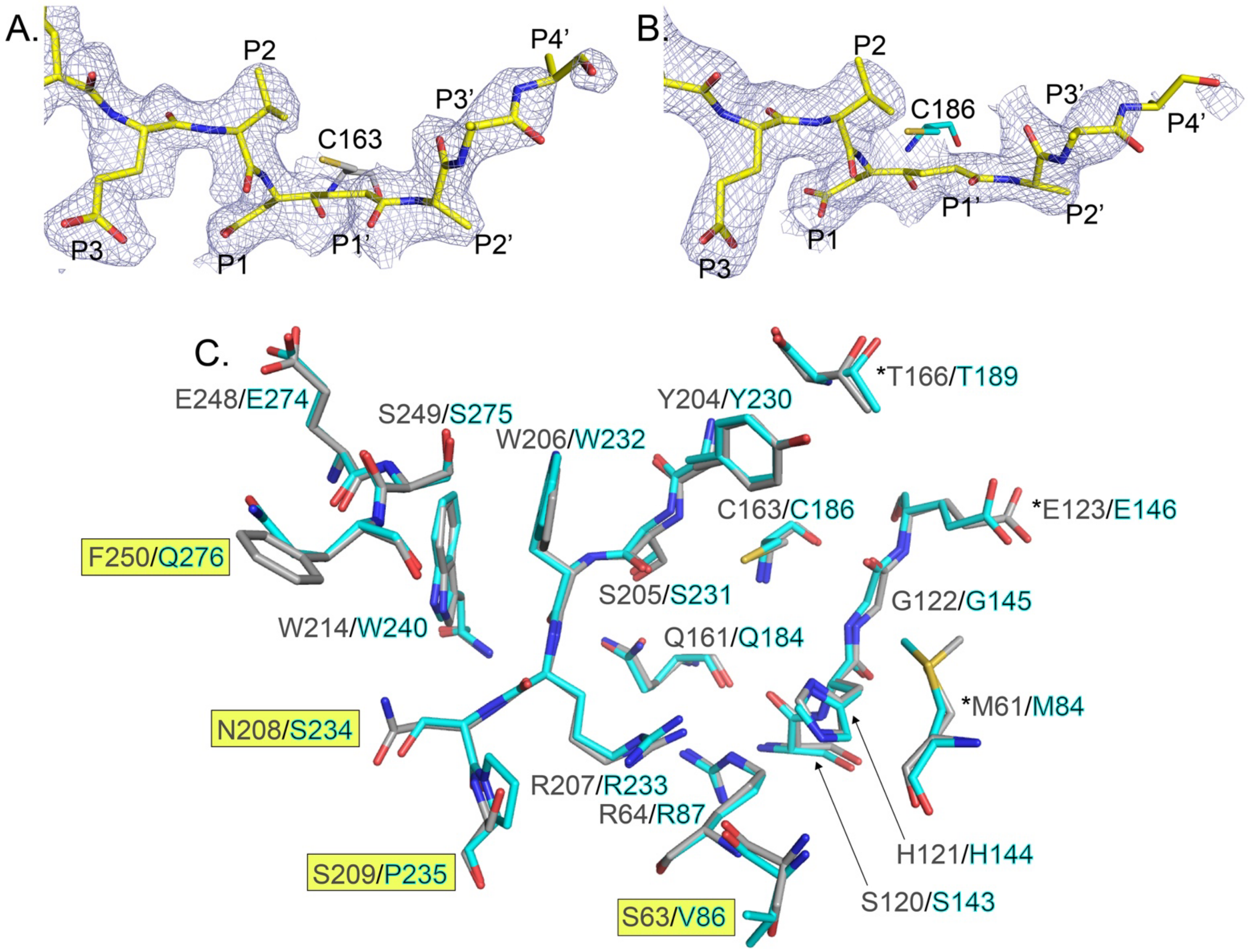


**Figure S4.** 2$f_o$-$f_c$ density contoured at 1σ of Ac-DEVD-Propionate-AAA bound to casp-3 (A) and casp-7 (B) showing highly resolvable density for P1’-P3’ bound to S1’-S3’ subsites. Casp-3 C163 and casp-7 C186 active-site cysteines are shown in grey and teal carbon, respectively. (C) Overlay of residues of casp-3 (grey carbon) and casp-7 (teal carbon) within 4 Å of Ac-DEVD-Propionate-AAA. Differences in amino acid side chains between the two active sites are highlighted by yellow box. * next to a pair denote these residues interact exclusively with the prime side of Ac-DEVD-Propionate.

**HPLC trace showing resolution of Ac-DEV(*D/L*)D(OBzl)-Propionate**

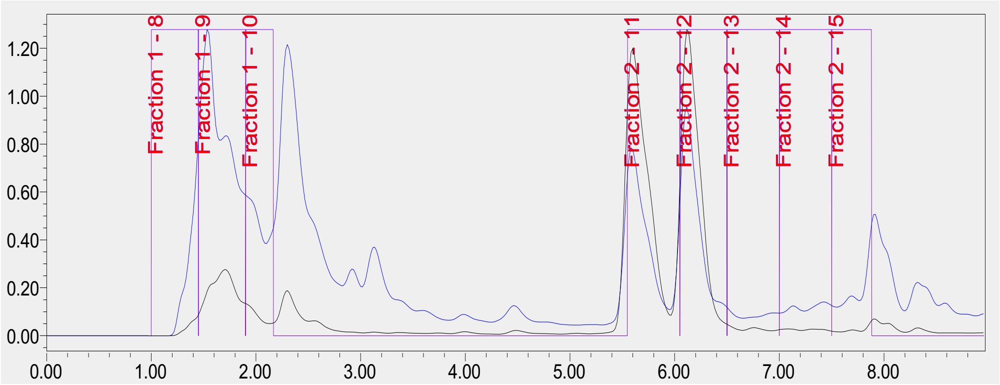


25 to 100% gradient over 13 min on 150 x 19 mm C18, 20 mL/min

**HPLC trace for Ac-DEV(*L*)D-Propionate (fraction 11 from above purification)**

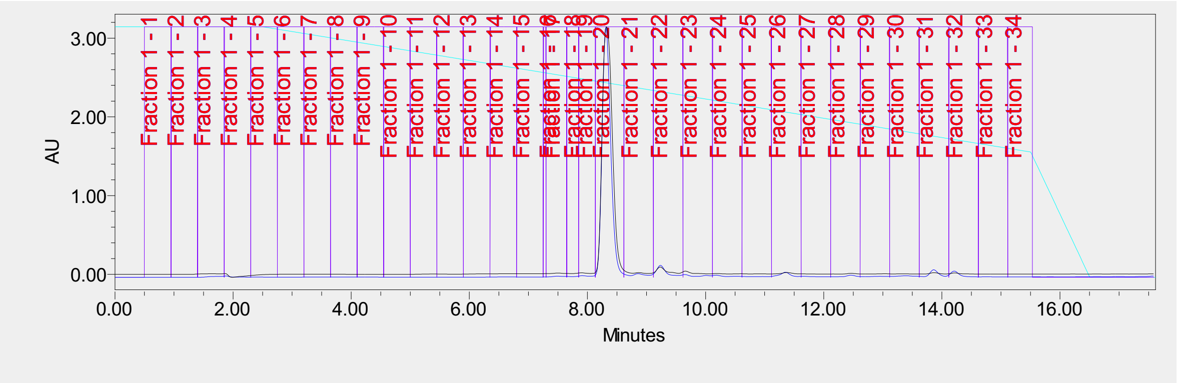


0 to 50% B over 13 min on 150 x 19 mm C18, 20 mL/min

**HPLC trace for Ac-DEV(*D*)D-Propionate (fraction 12 from above purification)**

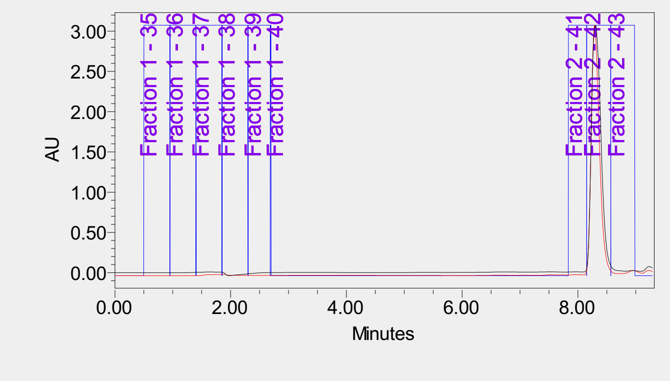


0 to 50% B over 13 min on 150 x 19 mm C18, 20 mL/min

**Table S1.** X-ray structure statistics for casp-8:Ac-DW3-KE structure

| | |
|---|---|
| PDB ID | 6X8H |
| Space Group | P $3_1$ 2 1 |
| a, b, c; Å | 62.5, 62.5, 129.4 |
| α, β, γ; ° | 90, 90, 120 |
| Resolution (Å) (outer shell) | 50.0-1.48 (1.51-1.48) |
| Completeness (%) | 99.8 (98.5) |
| Unique reflections | 49,543 (2,382) |
| Redundancy | 13.3 (7.0) |
| $R_{meas}$ (%)[a] | 13.3 (74.0) |
| $R_{merge}$ (%)[b] | 12.8 (68.4) |
| $R_{p.i.m.}$ (%)[c] | 3.6 (27.5) |
| Average I/Average σ (I) | 22.0 (2.8) |
| $CC_{1/2}$ | 97.0 (79.9) |
| Resolution (Å) (outer shell) | 50.0-1.48 (1.51-1.48) |
| No. reflections (test set)[d] | 49,494 (2,403) |
| $R_{cryst}$ (%)[e] | 14.9 (18.9) |
| $R_{free}$ (%) | 16.7 (22.3) |
| Protein atoms / Waters | 1,935 / 265 |
| CV coordinate error (Å)[f] | 0.11 |
| Rmsd bonds (Å) / angles (°) | 0.017 / 1.445 |
| B-values protein/waters/ligands ($Å^2$) | 12 / 28 / 14 |
| Ramachandran statistics (%) | |
| Preferred | 96.2 |
| Allowed | 3.8 |
| Outliers | 0 |

[a]$R_{meas} = \{\Sigma_{hkl}[N/(N-1)]1/2\Sigma_i|I_{i(hkl)} - \langle I_{(hkl)}\rangle|\} / \Sigma_{hkl}\Sigma_i I_{i(hkl)}$, where $I_{i(hkl)}$ are the observed intensities, $\langle I_{(hkl)}\rangle$ are the average intensities and N is the multiplicity of reflection hkl. [b]$R_{merge} = \Sigma_{hkl}\Sigma_i|I_{i(hkl)} - \langle I_{(hkl)}\rangle| / \Sigma_{hkl}\Sigma_i I_{i(hkl)}$ where $I_{i(hkl)}$ is the $i^{th}$ measurement of reflection h and $\langle I_{(hkl)}\rangle$ is the average measurement value. [c]$R_{p.i.m.}$ (precision-indicating $R_{merge}$) = $\Sigma_{hkl}[1/(N_{hkl} - 1)]^{1/2}\Sigma_i|I_{i(hkl)} - \langle I_{(hkl)}\rangle| / \Sigma_{hkl}\Sigma_i I_{i(hkl)}$. [d]Reflections with $I > 0$ were used for refinement[1-3]. [e]$R_{cryst} = \Sigma_h||F_{obs}|-|F_{calc}||/\Sigma|F_{obs}|$, where $F_{obs}$ and $F_{calc}$ are the calculated and observed structure factor amplitudes, respectively. $R_{free}$ is $R_{cryst}$ with 5.0% test set structure factors. [f]Cross-validated (CV) Luzzati coordinate errors.

**Table S2.** X-ray structure statistics for structures of casp-3 and casp-7 in complex with Ac-DEVD-Propionate

| Structure | Casp-3:Ac-DEVD-Propionate | Casp-7:Ac-DEVD-Propionate |
|---|---|---|
| PDB ID | 6X8I | 6X8J |
| Space Group | $P\ 2_1\ 2_1\ 2_1$ | $P\ 3_2\ 2\ 1$ |
| a, b, c; Å | 68.7, 85.0, 97.1 | 88.6, 88.6, 186.4 |
| α, β, γ; º | 90, 90, 90 | 90, 90, 120 |
| Resolution (Å) (outer shell) | 50.0-1.50 (1.53-1.50) | 50.0-2.60 (2.70-2.60) |
| Completeness (%) | 98.5 (99.4) | 99.8 (99.9) |
| Unique reflections | 90,329 (4,508) | 26,677 (1,273) |
| Redundancy | 5.0 (4.9) | 9.5 (9.6) |
| $R_{meas}$ (%)[a] | 7.3 (76.2) | 13.9 (141.7) |
| $R_{merge}$ (%)[b] | 6.5 (67.5) | 13.1 (134.1) |
| $R_{p.i.m.}$ (%)[c] | 3.2 (34.5) | 4.4 (45.1) |
| Average I/Average σ (I) | 21.4 (2.0) | 21.8 (2.1) |
| $CC_{1/2}$ | 96.3 (75.3) | 96.5 (87.6) |
| Resolution (Å) (outer shell) | 50.0-1.50 (1.52-1.50) | 50.0-2.60 (2.70-2.60) |
| No. reflections (test set)[d] | 90,209 (4,568) | 26,598 (1,333) |
| $R_{cryst}$ (%)[e] | 17.2 (23.2) | 18.3 (24.0) |
| $R_{free}$ (%) | 19.7 (25.8) | 21.1 (31.0) |
| Protein atoms / Waters | 3,791 / 355 | 3,670 / 43 |
| CV coordinate error (Å)[f] | 0.13 | 0.25 |
| Rmsd bonds (Å) / angles (º) | 0.016 / 1.463 | 0.003 / 0.609 |
| B-values protein/waters/ligands ($Å^2$) | 17 / 28 / 15 | 56 / 49 / 70 |
| Ramachandran statistics (%) | | |
| Preferred | 98.4 | 98 |
| Allowed | 1.6 | 2 |
| Outliers | 0 | 0 |

[a]$R_{meas} = \{\Sigma_{hkl}[N/(N-1)]1/2\Sigma_i|I_{i(hkl)} - <I_{(hkl)}>|\} / \Sigma_{hkl}\Sigma_i I_{i(hkl)}$, where $I_{i(hkl)}$ are the observed intensities, $<I_{(hkl)}>$ are the average intensities and N is the multiplicity of reflection hkl. [b]$R_{merge} = \Sigma_{hkl}\Sigma_i|I_{i(hkl)} - <I_{(hkl)}>| / \Sigma_{hkl}\Sigma_i I_{i(hkl)}$ where $I_{i(hkl)}$ is the $i^{th}$ measurement of reflection h and $< I_{(hkl)}>$ is the average measurement value. [c]$R_{p.i.m.}$ (precision-indicating $R_{merge}$) = $\Sigma_{hkl}[1/(N_{hkl} - 1)]^{1/2}\Sigma_i|I_{i(hkl)} - <I_{(hkl)}>|/\Sigma_{hkl}\Sigma_i I_{i(hkl)}$. [d]Reflections with I > 0 were used for refinement[1-3]. [e]$R_{cryst} = \Sigma_h||F_{obs}|-|F_{calc}||/\Sigma|F_{obs}|$, where $F_{obs}$ and $F_{calc}$ are the calculated and observed structure factor amplitudes, respectively. $R_{free}$ is $R_{cryst}$ with 5.0% test set structure factors. [f]Cross-validated (CV) Luzzati coordinate errors.

**Table S3.** X-ray structure statistics for structures of casp-3 and casp-7 in complex with Ac-DEVD-Propionate-AAA

| Structure | Casp-3:Ac-DEVD-Propionate-AAA | Casp-7:Ac-DEVD-Propionate-AAA |
|---|---|---|
| PDB ID | 6X8K | 6X8L |
| Space Group | $P\ 2_1\ 2_1\ 2$ | $P\ 3_2\ 2\ 1$ |
| a, b, c; Å | 69.9, 96.9, 100.5 | 88.3, 88.3, 187.3 |
| α, β, γ; ° | 90, 90, 90 | 90, 90, 120 |
| Resolution (Å) (outer shell) | 50.0-2.17 (2.21-2.17) | 50.0-2.45(2.49-2.45) |
| Completeness (%) | 97.5 (97.0) | 94.2 (94.7) |
| Unique reflections | 37,653 (1,845) | 56,614 (2,803) |
| Redundancy | 4.8 (4.1) | 2.9 (2.8) |
| $R_{meas}$ (%)[a] | 22.7 (154.0) | 13.8 (76.7) |
| $R_{merge}$ (%)[b] | 20.2 (134.9) | 11.4 (62.7) |
| $R_{p.i.m.}$ (%)[c] | 10.0 (72.3) | 7.6 (43.6) |
| Average I/Average σ (I) | 9.6 (2.1) | 9.4 (1.8) |
| $CC_{1/2}$ | 79.0 (25.1) | 94.1 (83.2) |
| Resolution (Å) (outer shell) | 50.0-2.17 (2.23-2.17) | 50.0-2.45 (2.53-2.45) |
| No. reflections (test set)[d] | 35,851 (1,833) | 30,516 (1,561) |
| $R_{cryst}$ (%)[e] | 22.0 (26.4) | 17.9 (25.3) |
| $R_{free}$ (%) | 26.3 (32.9) | 21.5 (30.7) |
| Protein atoms / Waters | 3,755 / 131 | 3,695 / 61 |
| CV coordinate error (Å)[f] | 0.28 | 0.29 |
| Rmsd bonds (Å) / angles (°) | 0.008 / 0.959 | 0.008 / 0.916 |
| B-values protein/waters/ligands (Å$^2$) | 28 / 30 / 34 | 50 / 47 / 68 |
| Ramachandran statistics (%) | | |
| Preferred | 98.4 | 98.4 |
| Allowed | 1.6 | 1.6 |
| Outliers | 0 | 0 |

[a]$R_{meas} = \{\Sigma_{hkl}[N/(N-1)]1/2\Sigma_i|I_{i(hkl)} - \langle I_{(hkl)}\rangle|\} / \Sigma_{hkl}\Sigma_i I_{i(hkl)}$, where $I_{i(hkl)}$ are the observed intensities, $\langle I_{(hkl)}\rangle$ are the average intensities and N is the multiplicity of reflection hkl. [b]$R_{merge} = \Sigma_{hkl}\Sigma_i|I_{i(hkl)} - \langle I_{(hkl)}\rangle| / \Sigma_{hkl}\Sigma_i I_{i(hkl)}$ where $I_{i(hkl)}$ is the $i^{th}$ measurement of reflection h and $\langle I_{(hkl)}\rangle$ is the average measurement value. [c]$R_{p.i.m.}$ (precision-indicating $R_{merge}$) = $\Sigma_{hkl}[1/(N_{hkl} - 1)]^{1/2}\Sigma_i|I_{i(hkl)} - \langle I_{(hkl)}\rangle| / \Sigma_{hkl}\Sigma_i I_{i(hkl)}$. [d]Reflections with $I > 0$ were used for refinement[1-3]. [e]$R_{cryst} = \Sigma_h||F_{obs}| - |F_{calc}|| / \Sigma|F_{obs}|$, where $F_{obs}$ and $F_{calc}$ are the calculated and observed structure factor amplitudes, respectively. $R_{free}$ is $R_{cryst}$ with 5.0% test set structure factors. [f]Cross-validated (CV) Luzzati coordinate errors.
